# Allele-specific correction of *ATAD3A* pathogenic variants via template-free CRISPR-Cas9 editing and gene conversion

**DOI:** 10.1101/2025.10.23.684255

**Authors:** Taejeong Bae, Yohan Park, Anel LaGrone, Milovan Suvakov, Ping Zhang, Hyekyung Park, Holly Van Remmen, James R. Lupski, Tamar Harel, Jean J. Kim, Alexej Abyzov, Wan Hee Yoon

**Affiliations:** Department of Quantitative Health Sciences, Center for Individualized Medicine, Mayo Clinic, Rochester, MN, USA; School of Biosystems and Biomedical Sciences, College of Health Science, Korea University, Seoul, Republic of Korea; Aging and Metabolism Research Program, Oklahoma Medical Research Foundation, Oklahoma City, OK, USA; Advanced Technology Cores, Human Stem Cell Core, Baylor College of Medicine, Houston, TX, USA; Oklahoma City VA Medical Center, Oklahoma City, OK, USA; Department of Molecular and Human Genetics, Baylor College of Medicine, Houston, TX, USA; Department of Pediatrics, Baylor College of Medicine, Houston, TX, USA; Human Genome Sequencing Center, Baylor College of Medicine, Houston, TX, USA; Texas Children’s Hospital, Houston, TX, USA; Department of Genetics, Hadassah Medical Center, Jerusalem, Israel; Faculty of Medicine, Hebrew University of Jerusalem, Israel; Stem Cells and Regenerative Medicine Center, Baylor College of Medicine, Houston, TX, USA

**Author notes:** These authors contributed equally to this work (Taejeong Bae, Yohan Park). **Corresponding authors:** Wan Hee Yoon, Ph.D., Aging & Metabolism Research Program, Oklahoma Medical Research Foundation, Oklahoma City, OK, 73104, USA, Alexej Abyzov, Ph.D. Department of Quantitative Health Sciences, Center for Individualized Medicine, Mayo Clinic, Rochester, MN 55905, USA.

**Keywords:** *ATAD3A*, allele-specific gRNA, template-free, CRISPR-Cas9, gene conversion, homologous recombination, dominant diseases, Harel-Yoon syndrome, HAYOS, mitochondria

## Abstract

Gene conversion is a specific form of homologous recombination (HR), involving the unidirectional transfer of genetic information from one genomic locus to another. CRISPR-Cas9-directed double strand breaks (DSBs) induce both interallelic and interlocus gene conversion in early human embryos and somatic cells, suggesting its potential for correcting pathogenic mutations. However, the key features in mitotic gene conversion, including its efficiency, the length of conversion track, and its dependency on specific recombination proteins, remain largely undefined. Here, we show that allele-specific CRISPR-Cas9-induced DSBs, without exogenous donor templates, can efficiently correct a heterozygous pathogenic variant (c.1582C>T; p.Arg528Trp) in the *ATAD3A* gene of patient-derived induced pluripotent stem cells (iPSCs). Amplicon-based next-generation sequencing (NGS) revealed that approximately 38%∼53% of edited iPSCs carried two wild-type *ATAD3A* alleles. Notably, over 99% of the corrected alleles derived from the homologous chromosome, indicating that the repair occurred mainly via interallelic gene conversion. Long-range amplicon nanopore sequencing coupled with haplotype analysis showed that the majority of gene conversion tracts was less than 2 kilobases in length. Whole-genome sequencing of three corrected iPSC clones showed the absence of large deletions or structural rearrangements at the *ATAD3A* target site. However, one clone carried a heterozygous deletion in *ATAD3B* locus, suggesting that CRISPR-Cas9 can introduce off-target genomic alterations. Knockdown of key HR proteins, including *RAD51*, *CtIP*, and *BRCA1/2*, significantly reduced the correction efficiency, indicating that the gene conversion relies on a RAD51-dependent HR pathway. Together, our findings provide compelling evidence that template-free CRISPR-Cas9-mediated interallelic gene conversion can be harnessed to correct disease causing variants in human iPSCs.

## INTRODUCTION

Gene conversion is a specific type of homologous recombination (HR) characterized by the unidirectional transfer of genetic information: the transfer of DNA from a donor sequence, typically a genomic region without a double-strand break (DSB), to an acceptor site where a DSB has occurred (Duret and Galtier 2009). The donor sequence can be interallelic (i.e., the other allele on the homologous chromosome) or nonallelic homologous sequences located at a different genomic locus (inter-locus) (Chen et al. 2007). During meiosis, gene conversion contributes to the exchange of genetic information between maternal and paternal DNA (Chen et al. 2007). Interlocus gene conversion, also referred to as non-allelic homologous recombination (NAHR), often occur between paralogous sequences, which contributes to sequence homogenization of these genes. However, this process increases the likelihood of genomic rearrangements, in some cases resulting in genetic disorders, such as Charcot-Marie-Tooth disease type 1A (CMT1A) and hereditary neuropathy with liability to pressure palsy (HNPP) (Inoue and Lupski 2002; Harel et al. 2016; Gunning et al. 2020). In contrast, mitotic gene conversion has been implicated in revertant mosaicism as observed in epidermolysis bullosa, a disorder caused by biallelic pathogenic mutations in the *collagen type XVII alpha 1 chain* (*COL17A1*) gene. In some patients, patches of clinically unaffected skin express full-length protein due to spontaneous gene conversion from wild-type sequence of the other allele, correcting the pathogenic variant in somatic cells (Jonkman et al. 1997; Jonkman and Pasmooij 2009). This example highlights the therapeutic potential of mitotic gene conversion as a form of natural gene therapy.

Mitotic gene conversion, a natural mechanism of somatic gene correction, provides a theoretical foundation for developing intentional gene conversion strategies in correcting pathogenic variants. Recent advances in genome editing, including Clustered Regularly Interspaced Short Palindromic Repeats (CRISPR)-Cas9, offer unprecedented opportunities to correct pathogenic mutations underlying genetic diseases (Jinek et al. 2012; Hsu et al. 2013; Doudna and Charpentier 2014). These tools now enable experimental testing whether intentional gene conversion can be harnessed for therapeutic use. In principle, this strategy can be implemented for both dominantly inherited diseases and recessively inherited diseases caused by compound heterozygous mutations. Recent studies have demonstrated that CRISPR-Cas9-mediated DSBs can result in the correction of pathogenic variants via gene conversion in human cells. For example, Mat et al. showed that CRISPR-Cas9 could correct a heterozygous pathogenic variant in the *MYBPC3* (*myosin binding protein C3*) gene, which causes hypertrophic cardiomyopathy (MIM: 600958), in human zygotes, suggesting that interallelic gene conversion can occur during early embryonic development in humans (Ma et al. 2017; Ma et al. 2018). In addition, CRISPR-Cas9 targeting the *hemoglobin subunit beta* (*HBB*) gene in HEK293T and human primary lung stromal cells resulted in interlocus gene conversion using the highly homologous *hemoglobin subunit delta* (*HBD*) as a template (Javidi-Parsijani et al. 2020). However, whether the approach can be extended to other somatic cell types, including induced pluripotent stem cells (iPSCs), remains unexplored. Moreover, detailed genomic analyses of CRISPR-Cas9-induced gene conversion events are still limited.

The ATPase family AAA-domain containing protein 3A (ATAD3A) is a mitochondrial membrane protein involved in mitochondrial DNA (mtDNA) organization and mitochondrial function (He et al. 2007; Gilquin et al. 2010). We and others have discovered that diverse genetic variations, including monoallelic or biallelic variants, deletion and duplication, and copy number variations (CNVs) in *ATAD3A* cause human neurodevelopmental disorders (NDD) (Harel et al. 2016; Desai et al. 2017; Gunning et al. 2020; Yap et al. 2021). To date, over 20 genetic variants in *ATAD3A* have been reported in association with NDD (Cooper et al. 2017; Desai et al. 2017; Peralta et al. 2019; Dorison et al. 2020; Hanes et al. 2020; Frazier et al. 2021). One such pathogenic allele is the recurrent *de novo* missense variant (NM_001170535.3; c.1582C>T: p.Arg528Trp) in exon 15 of *ATAD3A*. This heterozygous variant causes a distinct NDD, known as Harel-Yoon syndrome (HAYOS; MIM #617183), characterized by developmental delay, axonal neuropathy, optic atrophy, and/or hypertrophic cardiomyopathy (Harel et al. 2016). Using *Drosophila* and patient-derived fibroblasts, we previously demonstrated that the p.R528W variant acts as a dominant-negative mutation, leading to developmental lethality and severe mitochondrial dysfunction (Harel et al. 2016).

Humans have three *ATAD3* paralogs, *ATAD3A*, *ATAD3B*, and *ATAD3C,* arranged in tandem and sharing high sequence homology on chromosome 1p36.33. This arrangement likely arose from recently-evolved duplication of a single ancestral gene (Inoue and Lupski 2002; Li et al. 2014). Notably, we previously observed that CRISPR-Cas9-induced DSBs in exon 8 of *ATAD3A* can trigger interlocus gene conversion from *ATAD3B* to *ATAD3A* in HEK293T cells (Yanovsky-Dagan et al. 2022). Hence, intentional gene conversion by CRISPR/Cas9-mediated DSBs near the pathogenic site, could be a viable strategy for correcting *ATAD3A* pathogenic variants.

Here, we show that the delivery of a variant-specific Cas9 ribonucleoprotein (RNP) alone is sufficient to mediate efficient correction of the *de novo* pathogenic variant (c.1582C>T: p.Arg528Trp) in *ATAD3A* in human iPSCs derived from two unrelated individuals, even in the absence of an exogenous DNA template. Amplicon-based NGS revealed that over 99% of the corrected alleles originated from the homologous wild-type *ATAD3A* allele, indicating that the repair primarily occurred through interallelic gene conversion, rather than through interlocus events involving *ATAD3B*. Long-range amplicon nanopore sequencing and SNP haplotype analysis showed that the conversion track spanned less than 2 kbp. Moreover, knockdown of key HR factors, including *RAD51*, *CtIP/RBBP8, BRCA1,* and *BRCA2*, significantly reduced correction efficiency, indicating that RAD51-dependent HR is the primary mechanism driving interallelic gene conversion. Whole genome sequencing for three corrected iPSC lines confirmed the absence of large deletions and genomic rearrangements at the *ATAD3A* locus. However, one corrected clone exhibited a heterozygous deletion at *ATAD3B* locus, suggesting that CRISPR-Cas9 off-target activity can lead to unintended genomic alteration in regions of high sequence homology. In summary, our data provide compelling evidence that allele-specific CRISPR-Cas9 RNPs alone can induce template-free correction of pathogenic variants via gene conversion. However, our findings also reveal a critical limitation, as off-target activity of CRISPR-Cas9 at paralog loci can introduce unintended genomic alterations, underscoring the need for rigorous assessment of locus-specificity in genome editing experiments and the continued development of improved allele specificity of CRISPR-Cas9 designs to enhance both the safety and precision of genome editing strategies.

## RESULTS

### Gene corrected iPS cells without a donor template

In our previous study, we obtained skin fibroblasts carrying the *de novo* variant (c.1582C>T: p.R528W) in *ATAD3A* from individual II-2 (female) with HAYOS in family 1 (Harel et al. 2016). We also obtained peripheral blood mononuclear cells (PBMCs) carrying the same *de novo* variant in *ATAD3A* from an unrelated individual (female) presenting with HAYOS. Using Sendai virus-mediated reprogramming, we derived two iPSC clones from the fibroblasts of the first individual (C7 and C10) and two iPSC clones from the PBMCs of the second individual (C27 and C30) (**Figure 1A**). To assess the genomic integrity for the iPSC lines, we performed karyotyping by G-banding, which showed that all iPSC clones exhibited a normal karyotype (46, XX) (**Figure S1**). By performing flow cytometry analysis, we found that approximately 80-90% of all iPSC clones were positive for SSEA4 and OCT4, two hallmarks of human pluripotent stem cells (**Figure S2**).

**Figure 1.**
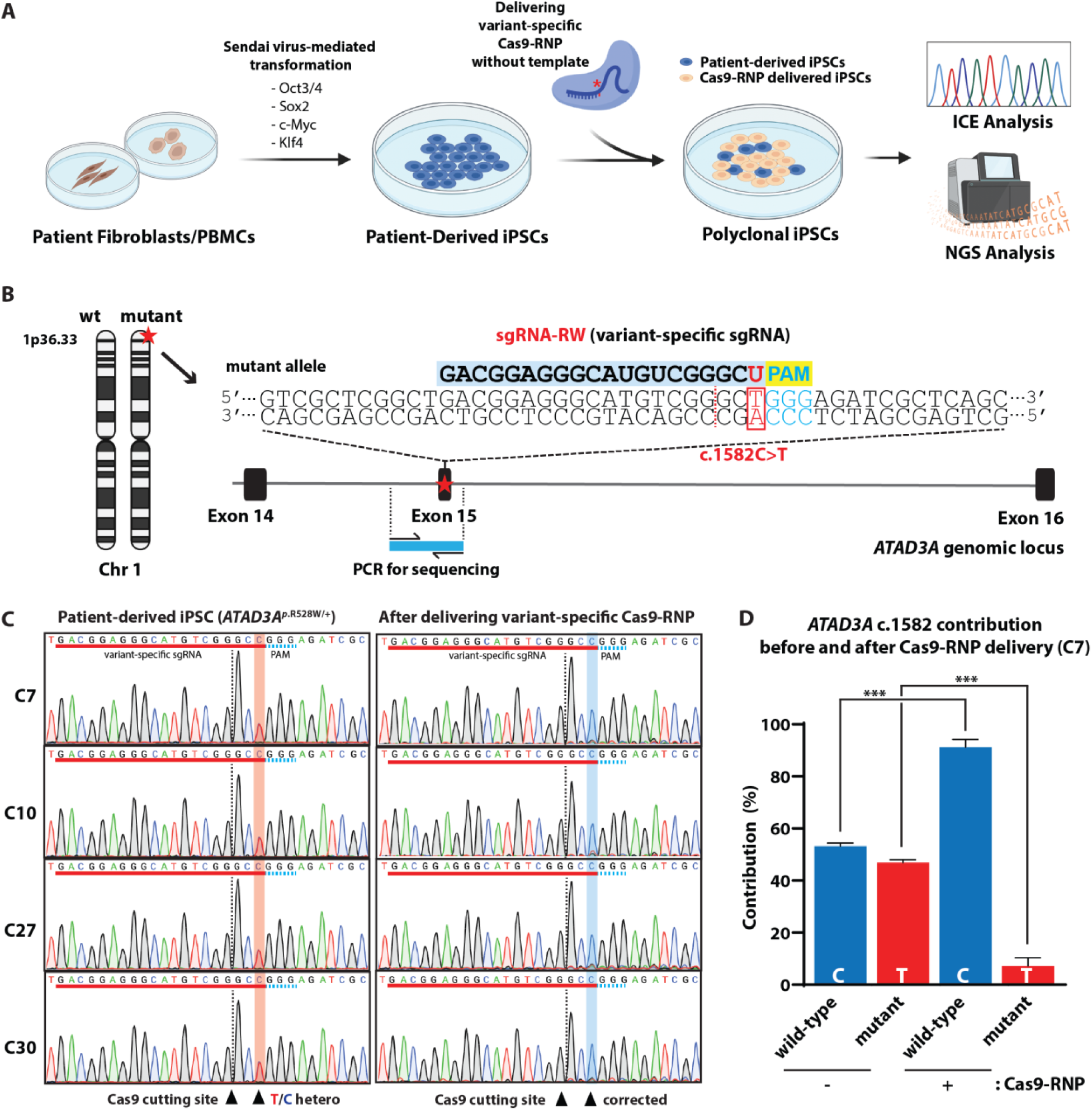
A delivery of variant-specific Cas9-RNP without a donor DNA template shows high-yield gene correction for the heterozygous *ATAD3A* variant. **(A)** A schematic for the generation of patient-derived iPSCs carrying the *de novo ATAD3A* variant (c.1582C>T: p.Arg528Trp) and delivery of the variant-specific Cas9-RNP without a donor template. **(B)** A schematic of the variant-specific sgRNA targeting the heterozygous variant (c.1582C>T) in exon 15 of the *ATAD3A* gene (sgRNA-RW). The variant-specific sgRNA is shown (blue color). The letter GGG (cyan) indicates PAM sequence. The blue bar indicates the genomic PCR region for subsequent genomic analysis. **(C)** Sanger chromatograms of iPSCs including C7, C10, C27, and C30 lines before and after the variant-specific Cas9-RNP delivery. Arrowheads with red box indicate the heterozygous c.1582C>T mutations in the patient-derived iPSCs. The blue boxes show their correction after the Cas9-RNP delivery. The arrowheads with dotted lines indicate Cas9 cutting sites. The red underlines show the sgRNA targeting sequences. The cyan dots indicate the PAM sequences. **(D)** ICE analysis of the Sanger chromatogram from the patient-derived iPSC (C7) after delivering the variant-specific Cas9-RNP shows an increase in C allele (91%) of the PCR products. Error bars indicate SEM (*n*=3). *P* values were calculated using Student’s t-test (\*\*\**P*<0.001).

To target the pathogenic variant allele (c.1582C>T) in *ATAD3A* in C7, we undertook a strategy employing CRISPR-Cas9-mediated mutant allele-specific genome editing (Smith et al. 2015; Burnight et al. 2017; Yamamoto et al. 2017; Giannelli et al. 2018; Gyorgy et al. 2019; Rabai et al. 2019). Cas9 endonuclease, including *Streptococcus pyogenes* Cas9 (*Sp*Cas9), was reported to tolerate mismatches between sgRNA and target DNA, but not those next to the protospacer adjacent motif (PAM) (Hsu et al. 2013). Hence, we designed a mutant allele-specific guide RNA in which the variant (c.1582C>T) is located next to the PAM sequence (sgRNA-RW) (**Figure 1B**). Cas9 was introduced into the iPSCs by electroporation of RNP consisting of sgRNAs and *Sp*Cas9 protein (**Figure 1A**) (Liang et al. 2015), which showed less off-site targeting compared to those in vector-mediated Cas9 overexpression (Kim et al. 2014). We electroporated the variant-specific Cas9 RNP (sgRNA-RW/*sp*Cas9) alone into the C7 iPSCs and then performed Sanger sequencing for the genomic PCR product (415 bps) around the target site (**Figure 1B**). Inference of CRISPR Editing (ICE) analysis (Conant et al. 2022) for the Sanger chromatogram of the genomic PCR product revealed ∼90% of the sequences were wild-type (C allele) in the edited C7 iPSCs (**Figure 1C and 1D**). We examined additional iPSC lines from the first individual (C10) as well as two iPSC lines (C27 and C30) from the second unrelated individual carrying the same *ATAD3A* variant (c.1582C>T: p.R528W). The electroporation-mediated delivery of the variant-specific Cas9 RNP led to an efficient T to C correction in all the iPSC lines (**Figure 1C**). Exon 15 of *ATAD3B* is a potential off-target site because it has 100% sequence identity with wild-type *ATAD3A* exon 15. The variant-specific gRNA is nearly complementary to the *ATAD3B* sequence, differing only in the last T base (**Figure S3**). Despite this high similarity, ICE analysis based on Sanger sequencing chromatograms revealed no evidence of indels in the ATAD3B region following delivery of the variant-specific Cas9 RNP (**Figure S4**). This finding suggests that the off-target editing at *ATAD3B* is minimum. Hence, these data show that the variant-specific targeting by CRISPR-Cas9 can correct the pathogenic c.1582C>T variant in *ATAD3A* without an exogenous DNA template.

### NGS for the PCR amplicon confirmed an efficient gene correction for the pathogenic *ATAD3A* variant

To assess sequence-level resolution of correction (C/T allele frequency) and indel frequencies, we performed next-generation sequencing (NGS) of the PCR amplicons from C7, C10, C27 and C30 lines before and after the Cas9 RNP delivery (**Figure 2A**). We found that the variant-specific Cas9 RNP (sgRNA-RW) delivery resulted in a significant increase in the C allele (65.2%∼71.8%) and a dramatic decrease in the T allele (6.7%∼13.3%) in all iPSC lines carrying the c.1582C>T variant (**Figure 2B**). In addition, the analysis revealed that the T allele has more indels than the C allele (**Figure 2B**), suggesting that the variant-specific Cas9 RNP preferentially targets the T allele. The corrected allele frequency suggests 38%∼53% of the iPSCs having both copies of the wildtype *ATAD3A* allele (c.1582C) after the Cas9 RNP delivery (**Table S1**). Hence, the NGS analyses confirmed that a template-free and variant-specific Cas9 RNP delivery can achieve correction of the *ATAD3A* p.R528W variant in multiple iPSC lines derived from unrelated families. In addition, the NGS analysis for *ATAD3B* amplicons showed a small increase in indel frequency with the C allele (0.9∼3.6%) in the four iPSC lines after delivering the variant-specific Cas9 RNP (**Table S2**), indicating that the variant-specific Cas9 RNP minimally targets wildtype *ATAD3B* allele.

**Figure 2.**
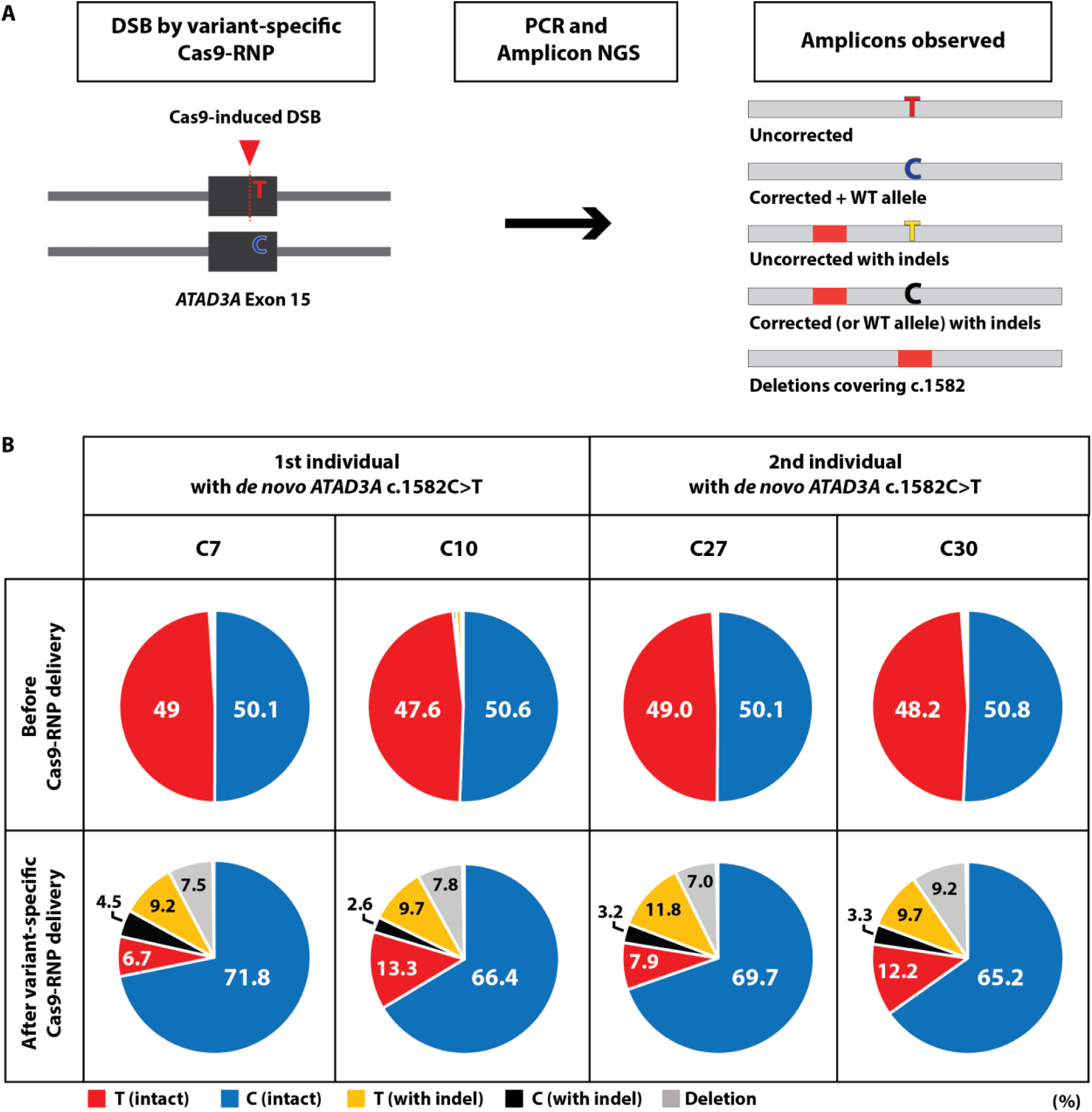
Amplicon NGS confirmed efficient gene correction of the heterozygous *ATAD3A* variant in iPSCs after the variant-specific RNP delivery. **(A)** A schematic of the heterozygous c.1582C>T variant in *ATAD3A* exon 15 and the variant-specific Cas9-RNP induced DSB (left) and observed outcomes (right). Five types of amplicons were observed; uncorrected c.1582 T allele, corrected/WT c.1582 C allele, uncorrected T allele with indels, corrected (or WT) C allele with indels, and deletions covering the c.1582 locus. **(B)** Amplicon NGS results show allele frequencies for *ATAD3A* c.1582 C allele (blue), *ATAD3A* c.1582 T allele (red), c.1582 C allele with indels (black), c.1582 T allele with indels (yellow), and deletions covering the c.1582 locus (grey) in C7, C10, C27, and C30 iPSC lines before and after the variant-specific Cas9-RNP delivery.

### High yield gene-correction in the absence of a template

To further confirm the gene correction of *ATAD3A* pathogenic variant using the template-free Cas9 approach, we performed subclonal selection and analysis on iPSCs to which the variant-specific Cas9-RNP had been delivered. We derived twenty clonal colonies from the C7 iPSC line after delivering the variant-specific RNP (**Figure 3A**). Sanger sequencing of genomic DNA of these clones showed that 70% of clones (14/20) carried only the wild-type allele, whereas 30% of clones (6/20) harbored indels around the cutting site of the sgRNA (**Figure 3B; Figure S5**). Higher percentage of corrected clones (70%) compared to the expected frequency of corrected cells (∼53%) from the NGS analysis (**Table S1**) suggests that the corrected clones outcompete uncorrected ones carrying the pathogenic variant during clonal selection.

**Figure 3.**
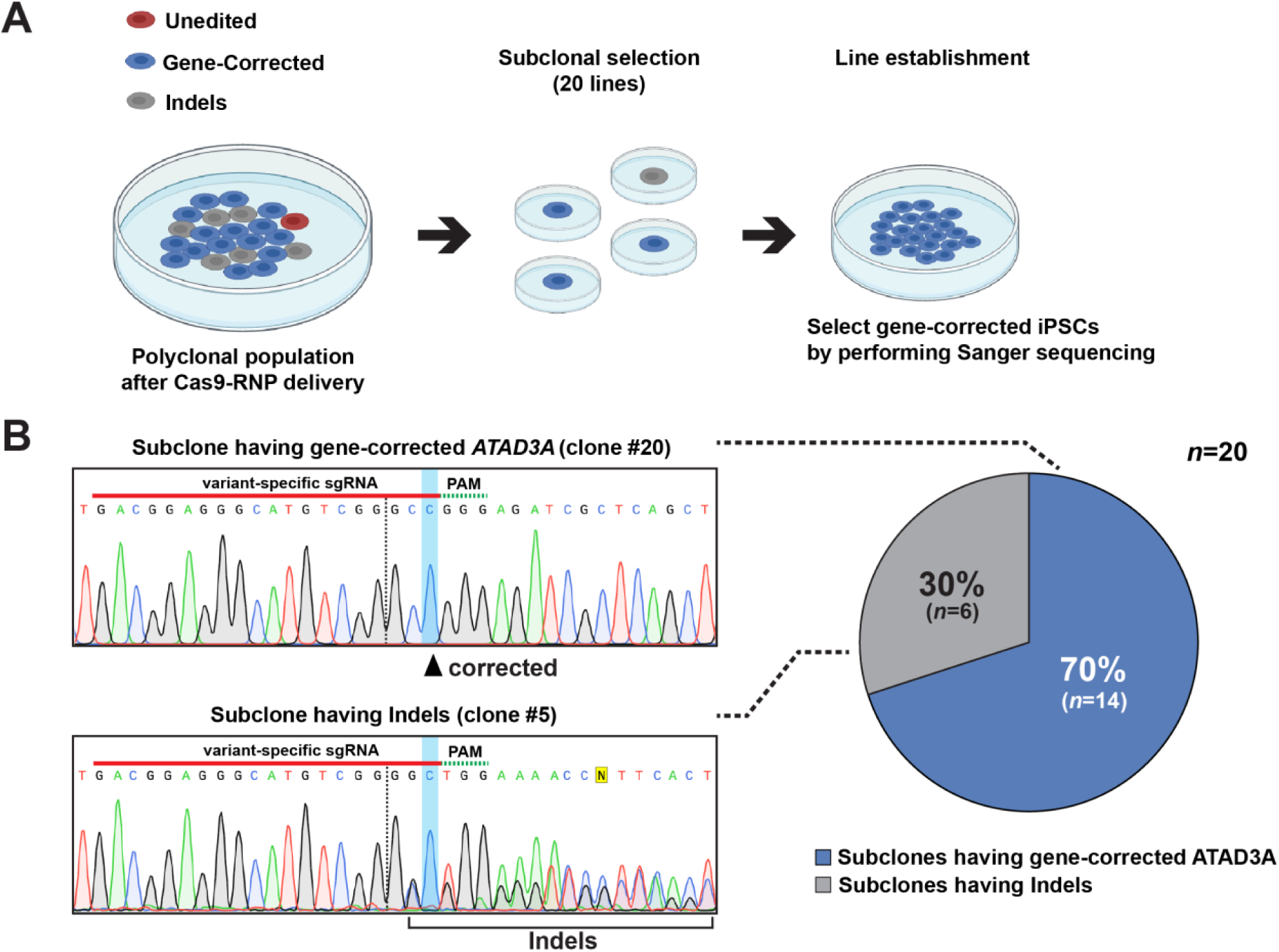
Subclonal selection of the variant-specific Cas9-RNP electroporated iPSCs shows high yield gene-correction in the absence of an exogenous template. **(A)** An illustration of subclonal selection after delivering the variant-specific Cas9-RNP to the patient-derived iPSCs (C7). Twenty subclones were selected and ICE analyses were performed to identify subclones with gene corrections. Additional NGS was performed to detect large indels in the *ATAD3* gene, copy number variation, as well as chromosomal integrity. **(B)** Representative Sanger chromatograms of gene-corrected and indel-harboring iPSC subclones (left). The arrowhead indicates a gene correction for the heterozygous *ATAD3A* variant (c.1582C>T) in subclone #20 (SC20). The presence of 1bp insertion resulted in overlapping sequencing peaks in subclone #5 (SC5). The red underlines indicate the target sequences of the variant-specific sgRNA. Cyan dots indicate PAM sequences. The pie chart shows the gene-correction efficiency of the variant-specific Cas9-RNP electroporation (right). 70% of subclones (*n*=14) showed the corrected variants (c.1582 C) in the *ATAD3A* gene (*n*=20).

### Interallelic gene conversion predominates in *ATAD3A* variant correction

Gene conversion can occur either between homologous alleles on the homologous chromosomes (interallelic) or between nonallelic homologous sequences located at a different genomic locus (interlocus) (Chen et al. 2007). In our previous study, we observed that in two out of nineteen sequenced HEK293T clones, CRISPR-Cas9-induced DSBs at exon 8 of *ATAD3A* led to interlocus gene conversion from *ATAD3B* to *ATAD3A* (Yanovsky-Dagan et al. 2022). To determine whether the correction of the c.1582C>T variant in *ATAD3A* resulted from interallelic or interlocus gene conversion, we analyzed *ATAD3A* amplicon sequences for the presence of *ATAD3B*-derived sequences before and after the Cas9 RNP delivery. Because *ATAD3A* and *ATAD3B* share a 222 bp region of identical sequence surrounding the target site, we quantified the nucleotide differences outside of this identical region to distinguish between the two genes (**Figure 4A**). Amplicon analysis revealed that only approximately 0.11% of *ATAD3A* reads contained *ATAD3B*-specific sequences, whereas the majority (>99.8%) corresponded to *ATAD3A* (**Figure 4B**). We also assessed whether reciprocal transfer of sequence information from *ATAD3A* to *ATAD3B* had occurred. Analysis of *ATAD3B* amplicons showed that reads containing *ATAD3A*-specific sequences accounted for less than 0.05% of total reads before and after Cas9 RNP delivery (**Figure 4B**), evidence showing no significant reciprocal exchange, indicating a unidirectional gene conversion mechanism. Collectively, these results demonstrate that the correction of DSBs at the *ATAD3A* c.1582T allele is predominantly mediated by interallelic gene conversion, with only a minor contribution (∼0.11%) from interlocus gene conversion.

**Figure 4.**
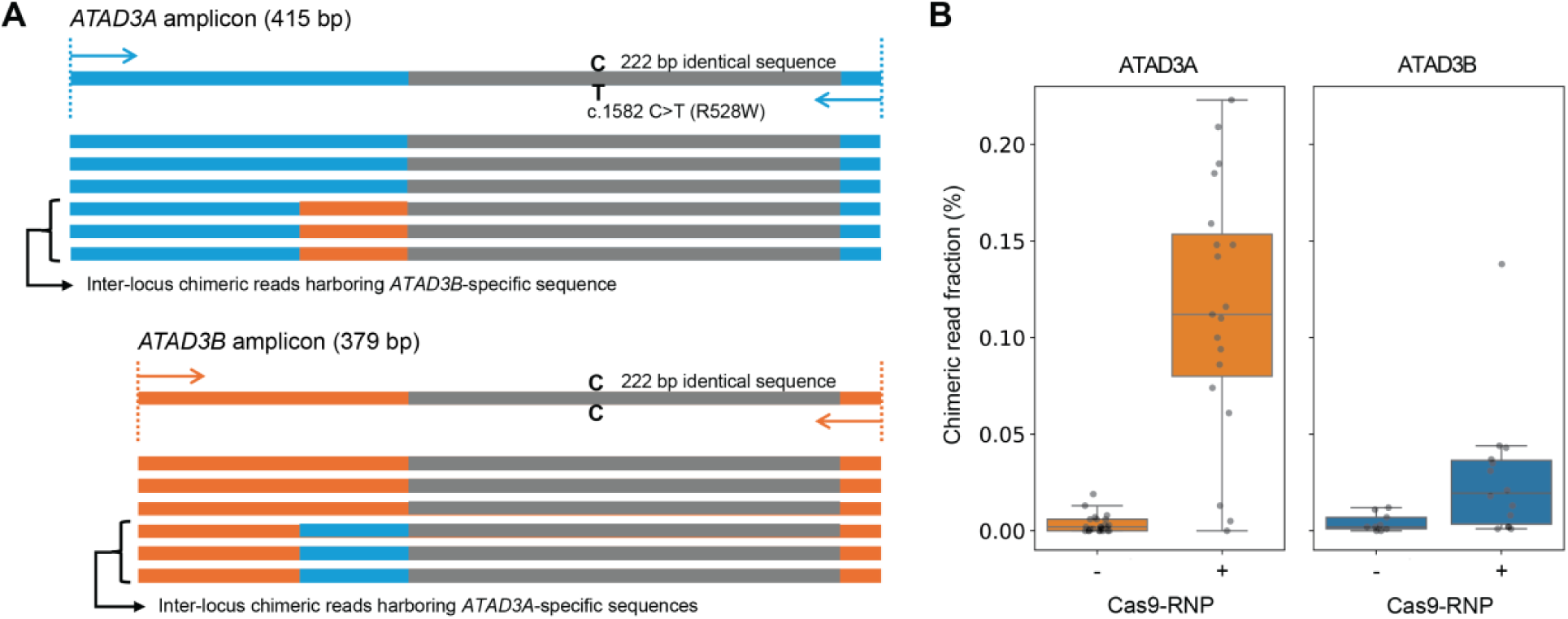
Interallelic gene conversion predominates in *ATAD3A* variant correction. **(A)** Schematic representation of inter-locus chimeric reads detected by amplicon NGS of *ATAD3A* and *ATAD3B* target regions. The *ATAD3A* amplicon (415 bp) carries the c.1582C>T variant, while the *ATAD3B* amplicon (379 bp) contains a 222-bp region identical to *ATAD3A* except for a T at the variant site. Chimeric reads were identified as those containing *ATAD3B*-specific sequences *ATAD3A* amplicons, or vice versa (Methods). The presence of such chimeric reads provides evidence that inter-locus gene conversion events occurred between the two paralogs. **(B)** Fraction of chimeric reads in *ATAD3A* and *ATAD3B* amplicon NGS samples before and after Cas9-RNP treatment. Cas9-RNP treatment increased chimeric reads, although overall fractions remained below 0.3%. The higher fraction in *ATAD3A* indicates that interlocus/interparalog gene conversion occurred predominantly from *ATAD3B* toward *ATAD3A*, providing a template for *ATAD3A* gene correction rather than in the opposite direction.

### Gene correction of p.R528W variant is primarily mediated by short-range gene conversion

Gene conversion can replace single-nucleotide variants (SNVs) flanking a DSB site by copying sequences from the homologous chromosome, often resulting in loss of heterozygosity (LOH)(Chen et al. 2007). Because unintended LOH can have detrimental cellular consequences, it is critical to determine the range of gene conversion track lengths when repairing DSBs generated by genome editing. Track length can be inferred by examining whether LOH occurs at SNPs surrounding the DSB site. Whole genome sequencing (WGS) of the C7 iPSC line identified three germline heterozygous SNPs near the c.1582C>T locus (chr1:1,463,318 C>A; chr1:1,465,382 A>G; and chr1:1,465,610 G>A) (**Figure 5A**). To monitor these SNPs, we performed Oxford Nanopore sequencing of 2,740 bps amplicons spanning the target site in C7 iPSCs, both before and after the Cas9 RNP delivery (**Figure 5A**). Only reads containing both flanking SNPs were retained for haplotype analysis. In unedited C7 iPSCs, we found that the majority (∼42.7%) of wild-type c.1582C (p.R528) alleles is associated with the haplotype C-C(p.R528)-A-G, whereas most pathogenic c.1582T (p.W528) alleles (∼41.4%) carried A-T(p.W528)-G-A (**Figure 5B and Table S3**). Interestingly, a small fraction (∼4.4%) of reads with the pathogenic haplotype had already undergone correction to A-C(p.R528)-G-A in the unedited C7 iPSCs, suggesting that spontaneous mitotic gene conversion can occur during iPSC culture. Another minor fraction (∼4.3%) displayed a chimeric haplotype, A-C(p.R528)-A-G, which contained SNP1 (chr1:1,463,318 A) from the pathogenic haplotype but carried the other wild-type c.1582C (p.R528) containing haplotype (**Figure 5B and Table S3**). The reciprocal haplotype, C-T(p.W528)-G-A, was detected at ∼3%, further supporting spontaneous recombination events during iPSC proliferation.

**Figure 5.**
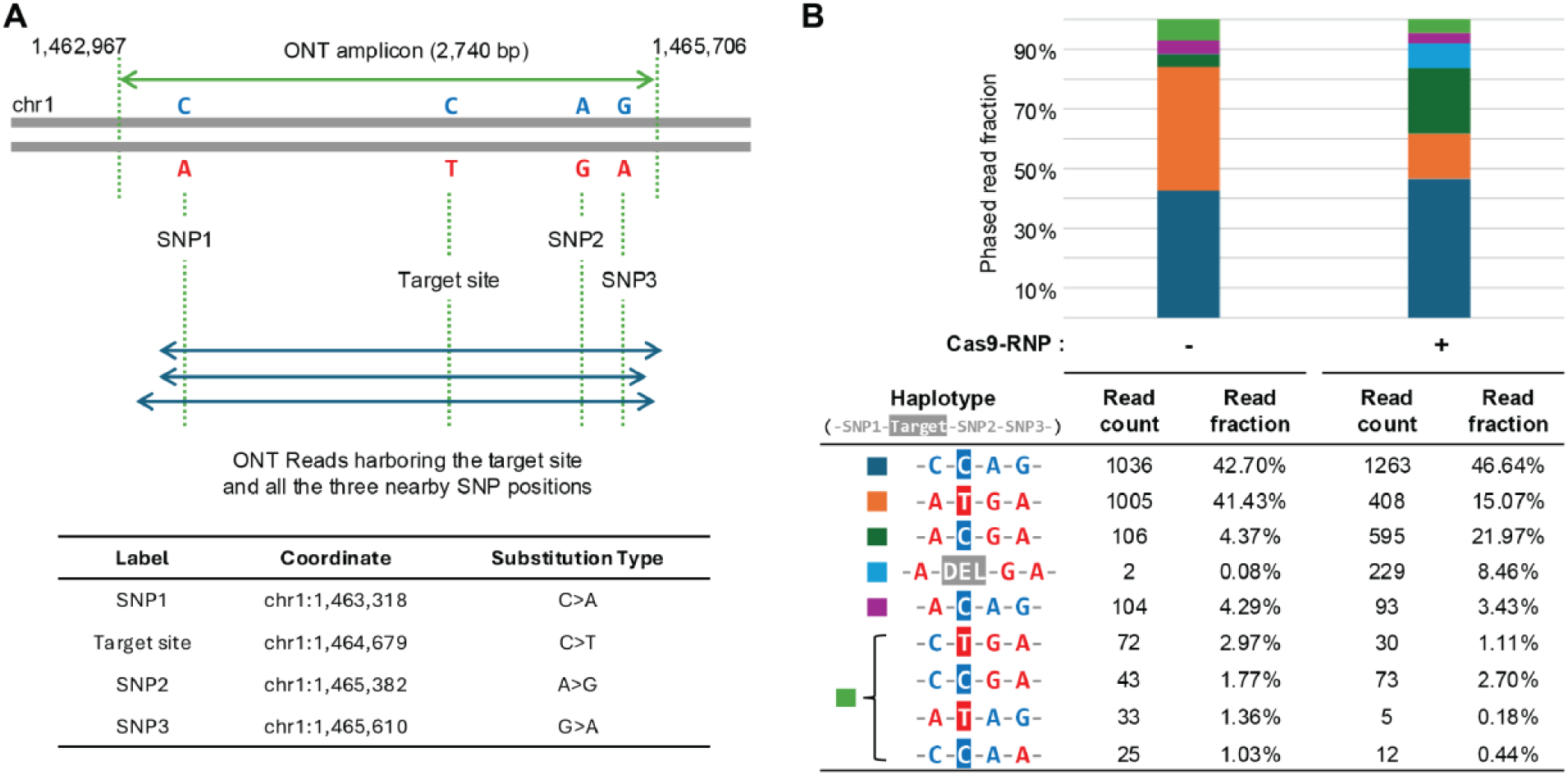
Gene correction of p.R528W is primarily mediated by short-range gene conversion. **(A)** Overview of the target region used for read-backed phasing. Long-range PCR amplified a 2,740-bp fragment encompassing the target site and three nearby SNPs, followed by long-read sequencing using Oxford Nanopore Technology (ONT). Only ONT reads containing the target variant and all three SNP positions were extracted and used for read-backed phasing. Genomic coordinates (GRCh37) and substitution types for these positions are listed below the schematic. **(B)** Fraction of phased reads of haplotypes in iPSCs before and after gene correction by Cas9-RNP. Each color in the stacked bar plot represents a distinct haplotype defined by the combination of alleles at SNP1, the target site, SNP2, and SNP3 (as shown in panel A). Eight haplotypes were identified as significant if their observed counts exceeded the expected frequency from haplotype error in either control or Cas9-RNP-treateds samples, as determined by a binomial test using haplotype error probability and total read counts (Method; see also Table S4). Four haplotypes with read fractions below 3% are grouped together and shown in a single color. Haplotypes with either the corrected reference allele (-A-C-G-A-) or a deletion (-A-DEL-G-A-) increased after Cas9-RNP treatment, with sequence changes limited to the target site, indicating short-range gene conversion between SNP1 and SNP2.

Following Cas9 RNP delivery, the corrected pathogenic haplotype (A-C(p.R528)-G-A) increased substantially to ∼22%, while the original pathogenic haplotype (A-T(p.W528)-G-A) decreased by ∼15.1% (**Figure 5B and Table S3**). The frequency of the wildtype haplotype C-C(p.R528)-A-G remained comparable to those in unedited control (∼42.7% vs ∼46.6%) and the proportion of the chimeric haplotype (A-C(p.R528)-A-G, ∼3.4%) remained similar to that of unedited cells (**Figure 5B and Table S3**). These results indicate that most Cas9-induced gene conversion events occur between SNP1 and SNP2 (chr1:1,463,318 C>A and chr1:1,465,382 A>G), spanning a region of 2,064 bp. In addition, Cas9 targeting of the c.1582T allele generated deletions near the target site (A-DEL-G-A) in ∼8% of the pathogenic haplotype (**Figure 5B and Table S3**). Collectively, these results reveal that the majority of gene conversion tracks are shorter than 2,064 bp, the distance between SNP1 and SNP2 and do not extend enough to cause LOH at these flanking SNPs.

### RAD51, BRCA1/2, and CtIP are required for efficient correction of *ATAD3A* pathogenic variant

Most HR requires RAD51, the ortholog of *E. coli* RecA, that plays a key role in strand invasion and DNA homology search (Prakash et al. 2015). However, RAD51-independent HR repair also has been reported, including intrachromosomal recombination, break-induced recombination (BIR), single-strand template repair in *S. cerevisiae* (Bai and Symington 1996; Ira and Haber 2002; Gallagher et al. 2020) and break-induced telomere synthesis in human cell lines (Dilley et al. 2016). Notably, CRISPR-Cas9-mediated single-strand template repair does not require RAD51 in human cells (Richardson et al. 2018). To determine whether the gene correction of the pathogenic *ATAD3A* variant by the CRISPR-Cas9 editing requires RAD51, we performed siRNA-mediated knockdown for *RAD51* in the C7 iPSCs (**Figure 6A**) when inducing DSBs in the c.1582C>T (p.R528W) allele by the Cas9 RNP. NGS analysis showed that *RAD51* knockdown leads to a significant reduction of C allele correction and increased T allele frequency (**Figure 6B**). Hence, the template-free and variant-specific *ATAD3A* gene correction requires RAD51.

**Figure 6.**
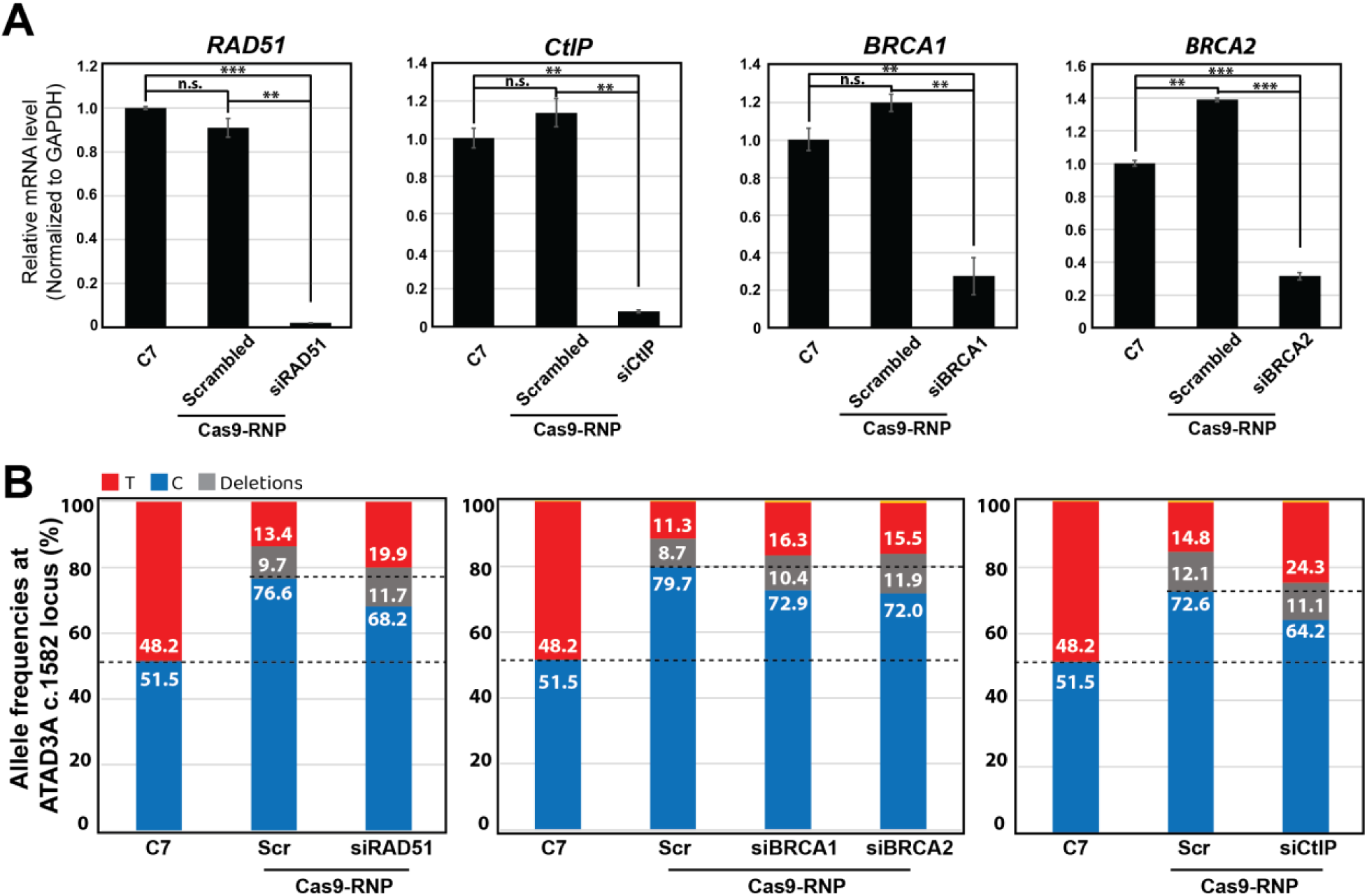
RAD51, BRCA1, BRCA2, and CtIP are required for the template-free gene correction in *ATAD3A* c.1582C>T variant. **(A)** Quantitative RT-PCR results show the mRNA levels of *RAD51*, *CtIP*, *BRCA1*, and *BRCA2* in C7 iPSCs following siRNA treatment. Error bars indicate SEM. *P* values were calculated using Student’s t-test. \*\**P*<0.01, \*\*\**P*<0.001. N.S. indicates not statistically significant. **(B)** Amplicon NGS reveals allele frequencies of *ATAD3A* c.1582 locus after the variant-specific Cas9-RNP with or without siRNAs targeting *RAD51*, *BRCA1*, *BRCA2*, and *CtIP*.

Next, we questioned if *BRCA1* (*BRCA1 DNA repair associated*) and *BRCA2* (*BRCA2 DNA repair associated*) are required for *ATAD3A* gene correction, since both proteins play a key role in recruiting RAD51 to DSBs (Prakash et al. 2015). BRCA1 plays a role in two distinct steps: 5’ to 3’ resection of DSBs to generate 3’ ssDNA overhangs by directly interacting with the resection factor CtIP (Wong et al. 1998; Yu et al. 1998) and loading of the RAD51 recombinase into the ssDNA through interacting with PALB2-BRCA2 (Xia et al. 2007; Sy et al. 2009). We sought to determine whether BRCA1, BRCA2, and/or CtIP are required for correcting the *ATAD3A* pathogenic variant by CRISPR-Cas9-induced gene conversion. NGS analyses showed that knockdown of all three proteins negatively impact gene correction (**Figure 6A and 6B**). Hence, the results indicate that the BRCA1 and CtIP-mediated end resection and BRCA1/2-mediated RAD51 loading are required for the *ATAD3A* gene correction.

### Genomic Analyses for on-target and off-target indels and genomic structural integrity

The major on-target effects of CRISPR-Cas9 are thought to be small indels of less than 20 base pairs (bps) (Koike-Yusa et al. 2014; Tan et al. 2015; van Overbeek et al. 2016). CRISPR-Cas9-directed genome editing, however, was also reported to cause large deletions (kilobase-scale) and/or complex genomic rearrangements at the targeted sites in mouse and human cells (Kosicki et al. 2018). Thus, genotyping by performing PCR that captures only a small genomic region centered on the Cas9 target site will not determine whether the gene editing resulted in larger deletion or structural rearrangements of DNA that remove primer-binding sites, leading to amplification of only the wild-type allele. To assess whether CRISPR-Cas9-directed DSBs could lead to large deletions at the target site in the *ATAD3A* genomic locus, we performed whole genome sequencing (WGS) at 30x coverage on the patient-derived iPSC (C7) and three gene-corrected iPSC lines (SC20, SC1, and SC14) (**Figure 7A**). WGS revealed no evidence of large deletions in the nearby genomic region surrounding the Cas9 target site in *ATAD3A* exon 15 of SC20, SC1, or SC14 cells (**Figure 7A**). In addition, all three corrected iPSC clones carried heterozygous SNPs (chr1:1,463,318 C>A; chr1:1,465,382 A>G; and chr1:1,465,610 G>A), indicating that short-range (∼2 kbp) gene conversion mediated the correction of the pathogenic variant. However, structural analysis revealed that while SC20 and SC1 maintained an intact genomic structure in the *ATAD3B* region, SC14 clone carried a heterozygous deletion of approximately 40 kb spanning from *ATAD3B* to *ATAD3A* (**Figure 7B and Figure S6**). Given that NAHR between *ATAD3A* and *ATAD3B* has been reported to occur during meiosis (Harel et al. 2016), this deletion is likely the result of NAHR between these paralogous genes during the DNA repair process.

**Figure 7.**
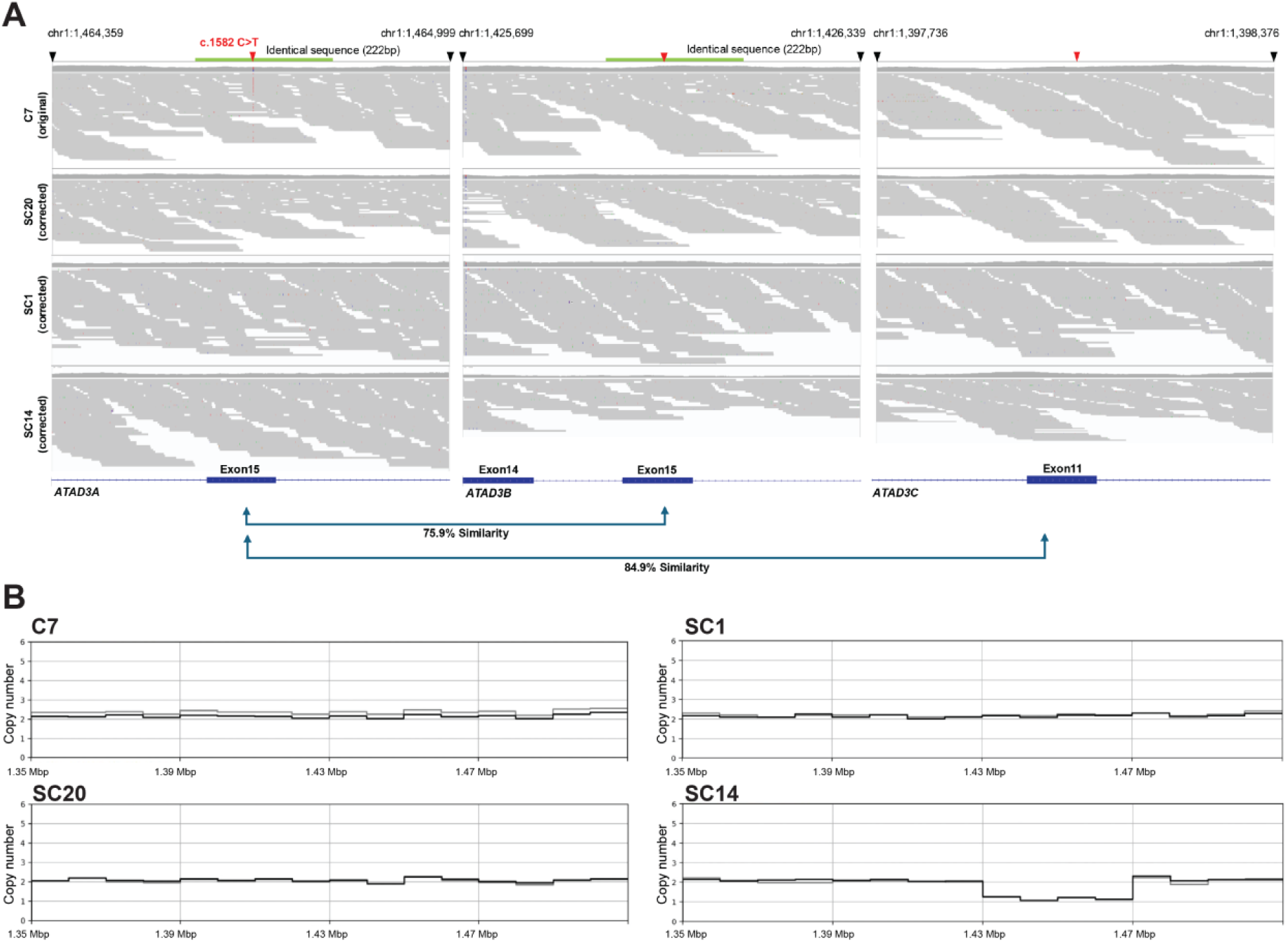
Genomic Analyses for on-target and off-target indels and genomic structural integrity. **(A)** Integrative Genomics Viewer (IGV) snapshots showing *ATAD3A* exon 15 region and the homologous regions in *ATAD3B* and *ATAD3C* in the patient-derived iPSC clone (C7) and three gene-corrected iPSC clones (SC20, SC1, and SC14). Red arrowheads indicate the c.1582C>T variant and corresponding loci in the paralogs. Green bars mark the boundaries of the 222-bp identical sequences shared between *ATAD3A* and *ATAD3B*. **(B)** Copy number profiles across chromosome 1 spanning *ATAD3A*, *ATAD3B* and *ATAD3C*, represented by read depth Manhattan plots (bin size = 10 kb) in the same iPSC lines (C7, SC20, SC1, and SC14). A deletion between the immediate upstream regions of *ATAD3B* exon 15 and *ATAD3A* exon 15 was detected in the SC14 clone (see also Figure S6).

### Restored mitochondrial function in the corrected iPSC clone

We sought to determine cellular function associated with mitochondria in the corrected iPSC clones, as ATAD3A dysfunction was shown to negatively affects mitochondrial respiration (Jin et al. 2018). To determine the mitochondrial respiration in both patient (C7) and the gene corrected iPSC clones (SC20), we performed a Seahorse metabolic assay that measures cellular respiration and mitochondrial function (Pharaoh et al. 2019). We found that the patient iPSCs (C7) exhibited a decrease in both the basal oxygen consumption rate (OCR) and the maximal oxidative capacity compared to those in the gene-corrected iPSCs (SC20) (**Figure 8A, 8C and 8D**). Accordingly, we found that the ATP-linked respiration is significantly lower in the patient iPSCs (C7) than that in the gene-corrected iPSCs (SC20) (**Figure 8E**). Hence, these results demonstrate that the heterozygous p.R528W variant leads to defects in mitochondrial respiration, which was rescued by the gene correction for the p.R528W variant in *ATAD3A*.

**Figure 8.**
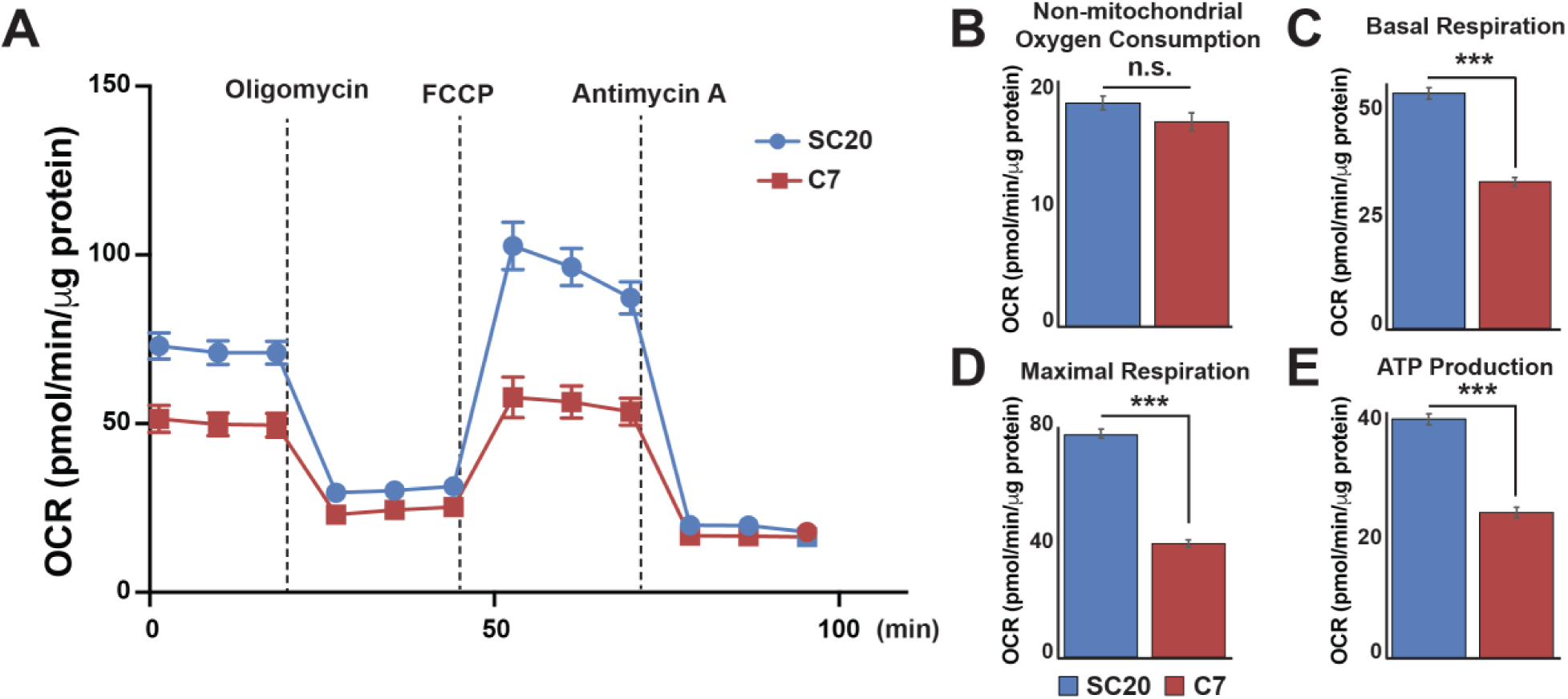
Correction of the ATAD3A pathogenic variant restores mitochondrial respiration. Oxygen consumption rates were measure in patient-derived iPSCs (C7) and gene-corrected (SC20) iPSC lines. Data represent nine biological replicates per cell line. Error bars indicate SEM. *P* values were calculated using Student’s t-test (\*\*\**P*<0.001. N.S. indicates not statistically significant).

## DISCUSSION

DSBs induced by genome editing in somatic cells are predominantly repaired thorough the error-prone non-homologous end joining (NHEJ) pathway, often resulting in insertions or deletions (indels) at the target sites (Lieber 2010). In contrast, the homology-directed repair (HDR) pathway is engaged less frequently and requires the presence of a homologous DNA template (Jackson and Bartek 2009). Hence, gene correction typically requires co-delivery of CRISPR-Cas9 and an exogenous homologous DNA template (Smith et al. 2015; Rabai et al. 2019). In contrast to these prior findings, we demonstrate here efficient gene correction of the recurrent *de novo ATAD3A* variant (c.1582C>T, p.R528W) in iPSCs using allele-specific Cas9 RNP without the need for an exogeneous DNA template. Amplicon NGS for the patient iPSCs after the RNP delivery revealed that 65∼70% of the sequences were wild type. Mechanistically, we found that the gene correction process primarily relies on the canonical gene conversion events, including end resection and a RAD51-dependent homologous recombination pathway.

Gene conversion can occur either between homologous alleles located on the homologous chromosomes (interallelic) or between nonallelic homologous sequences located at a different genomic locus (interlocus) (Chen et al. 2007). Initially, we hypothesized that the gene correction in the iPSCs might result from NAHR (interlocus gene conversion) between *ATAD3A* and its paralog *ATAD3B*. This hypothesis was based on the tandem arrangement of high sequence homology between these paralogs, which renders this genomic region prone to NAHR during meiosis (Carvalho and Lupski 2016; Harel et al. 2016; Gunning et al. 2020; Yap et al. 2021). Furthermore, our previous CRISPR-Cas9 experiments in HEK293T cells revealed that DSBs induced in *ATAD3A* led to gene conversion events in 2 out of 19 sequenced clones, with *ATAD3B* used as a template (Yanovsky-Dagan et al. 2022). However, our amplicon sequencing analysis showed that only 0.11% of *ATAD3A* reads contained *ATAD3B*-specific sequences, whereas the majority (>99.8%) of the reads corresponded to *ATAD3A* (**Figure 4B**), indicating that the predominant mechanism of gene conversion is interallelic gene conversion. This observation aligns with prior studies on the *HBB* gene, where CRISPR-Cas9-induced DBSs in the *HBB* gene resulted in only 0.15% of interlocus gene conversion events using *HBD* as a donor in human primary cells (Javidi-Parsijani et al. 2020). Given that *ATAD3A* and *ATAD3B* share a 222 bp region of identical sequence surrounding the target site, we cannot entirely rule out the possibility of short-track interlocus gene conversion occurring within this identical 222 bp DNA.

Interallelic gene conversion involves replacing sequences from a homologous chromosome having SNPs or variants nearby a DSB site, often resulting in LOH. LOH can be local, spanning only a few hundreds of bps to a few kbps. However, it can extend much further when driven by crossover events or break-induced replication (Saini et al. 2013; Mayle et al. 2015). This phenomenon is particularly related to cancer biology, where LOH contributes to the progression of tumors harboring recessive mutations in *RB1* or *TP53* (LaRocque et al. 2011). Hence, in genome editing applications, including those involving CRISPR-Cas9, understanding the typical range of gene conversion tracks is crucial for minimizing unintended LOH. Our nanopore sequencing of a 2,740 bp amplicon containing three SNP revealed that most correction events (∼22%) did not induce LOH across all three SNPs. In addition, all three corrected iPSC clones carried heterozygous SNPs (chr1:1,463,318 C>A; chr1:1,465,382 A>G; and chr1:1,465,610 G>A). This shows that gene conversion tracks generally span within 2,064 bp, the distance between first two SNPs, without extending far enough to disrupt genomic integrity. Interestingly, even in unedited iPSCs, we observed low frequency SNP switching, suggesting that spontaneous gene conversion may occur during normal cell cycle.

While the most on-target effects of CRISPR-Cas9 are typically small indels (Koike-Yusa et al. 2014; Tan et al. 2015; van Overbeek et al. 2016), accumulating evidence has shown that CRISPR-Cas9 can also induce large-scale deletions and/or genomic rearrangements in mouse and human cells (Kosicki et al. 2018). Conventional PCR-based genotyping that amplifies a short region (450 bp to a few kbp) centered on Cas9 target site may fail to detect large deletion or rearrangements. To address this limitation, we performed WGS at 30x coverage on the patient-derived iPSC line (C7) and three iPSC clones (SC20, SC1, and SC14). While no large deletions were detected at the *ATAD3A* exon 15 target stie in any of the edited lines, structural analysis uncovered a notable exception: SC14 harbored a heterozygous ∼40 kbp deletion spanning from *ATAD3B* exon15 to a region 3 kbp upstream of *ATAD3A* exon 15. In contrast, SC20 and SC1 retained intact genomic structure across *ATAD3B* and *ATAD3A* region. Given the off-target activity (∼0.4∼1.1%) of the allele-specific sgRNA/Cas9 toward the wildtype C allele in *ATAD3B* (**Table S2**) and prior reports of NAHR between these paralogs during meiosis (Harel et al. 2016), it is plausible that the deletion in SC14 resulted from NAHR triggered during the DNA repair process for DSBs in *ATAD3B* exon 15. This finding emphasizes the importance of comprehensive genomic analysis when evaluating CRISPR-Cas9 editing outcomes, especially in regions with paralogous sequences. Moreover, improving the allele specificity of sgRNA/Cas9 designs will be essential to minimize unintended recombination events and enhance the precision of genome editing.

Inference of CRISPR Editing (ICE) and TIDE analysis are widely used tools for analyzing CRISPR-Cas9 editing outcomes based on Sanger chromatogram (Conant et al. 2022). However, we observed that ICE analysis tends to overestimate the frequency of corrected C alleles in edited samples (**Figure 1**), compared to results obtained from amplicon-based NGS (**Figure 2**). This discrepancy shows a limitation of Sanger-based inference methods. Therefore, when precise quantification of sequence correction and indel frequencies is required, NGS is the more reliable approach for assessing accurate, sequence-level resolution of correction and indel reads.

Since our initial finding of the *de novo ATAD3A* p.R528W variant as a pathogenic mutation (Harel et al. 2016), numerous pathogenic alleles at the locus, including both monoallelic and biallelic variants in *ATAD3A* have been described (Cooper et al. 2017; Desai et al. 2017; Peralta et al. 2019; Dorison et al. 2020; Gunning et al. 2020; Hanes et al. 2020; Frazier et al. 2021; Yap et al. 2021). *ATAD3A* now appears to be the most common gene locus that results in lethal neonatal mitochondrial disease (Frazier et al. 2021). Recently, we found that missense variants or single nucleotide variants (SNVs) *in trans* to deletion or frameshift alleles lead to varied severity of phenotypes ranging from neonatal lethality to hypotonia, global developmental delay, learning difficulties, and ataxia (Yap et al. 2021). Adult heterozygous carriers who harbor one copy of *ATAD3A* loss-of-function alleles, however, exhibited no substantial health problems, indicating that one intact copy is sufficient for normal human development and physiology (Harel et al. 2016; Yap et al. 2021). Hence, gene correction of one pathogenic missense or SNV allele would be anticipated to be sufficient for restoring biological balance for individuals with biallelic variants in *ATAD3A.* The approach for the gene correction of *ATAD3A* variants reported in this study offers a promising framework for targeting additional pathogenic variants in *ATAD3A*.

In this study, we show that the delivery of a mutant allele-specific sgRNA/*Sp*Cas9 RNP is an effective way for gene correction with the heterozygous pathogenic variant in *ATAD3A* in iPSCs. It will be important to further explore this method for the other types of human cells as well as pathogenic variants in the genes having highly homologous paralogs (e.g., *SMN1*and *SMN2*) to achieve gene correction by intentional gene conversion.

## MATERIALS AND METHODS

### Study Design

The objective of this study was to determine whether intentional gene conversion, induced by CRISPR-Cas9, could correct pathogenic variants in *ATAD3A* in patient-derived iPSCs, and to determine the underlying molecular mechanism. This study used iPSCs derived from individuals diagnosed with HAYOS who carry pathogenic variants in *ATAD3A*. We measured the allele frequency of pathogenic allele both before and after delivering variant-specific Cas9 RNP, by performing ICE analysis from Sanger chromatograms and performing NGS analyses for PCR amplicons of target sites. The numbers of replications varied and is specified in each figure. The study was not blinded.

### Cell culture and iPSC generation

This study was approved by an institutional review board (IRB) at Oklahoma Medical Research Foundation, and written informed consent was obtained prior to genetic testing and sample collection. Primary fibroblasts from patient (individual II-2 in family 1) with the *ATAD3A* c.1582C>T, p.R528W*/+* variant were obtained in a previous study (Harel et al. 2016). PBMCs carrying the same *de novo ATAD3A* variant (c.1582C>T) were collected from individuals with HAYOS. Patient fibroblasts and PBMCs were reprogrammed by using CytoTune^TM^ iPS 2.0 Sendai Reprogramming Kit (Thermo Fisher # A16517) according to the manufacturer’s guidelines. Briefly, 2.0 x 10^5^ fibroblasts/PBMCs were transduced with a Sendai virus cocktail encoding hOct3/4, hSox2, hc-Myc and hKlf4. The virus cocktail was removed after 24 hours. After 5 days, cells were detached with Accutase (StemCell Technologies, Cat # 07920) and seeded onto hESC-qualified Matrigel (Corning Cat # 354277)-coated 10-cm culture plates. After 21 days, iPSC colonies were picked and plated onto Matrigel-coated 24-well plates. Patient-derived iPSCs and gene-corrected iPSCs were cultured in mTeSR1 medium (StemCell Tech # 85850) in Matrigel-coated plates at 37°C in a 5% CO_2_ humidified incubator.

### CRISPR/Cas9 Design, Electroporation for Cas9-RNP

The variant-specific gRNA was designed with help of Benchling’s prediction program as well as TrueGuide^TM^ gRNA design service (Thermo Fisher Scientific) (Table S4). The sgRNA carrying the variant-specific gRNA was synthesized by Synthego (Redwood, California). For gene editing, *Sp*Cas9-sgRNA RNP complexes were delivered into iPSCs using Neon^TM^ Transfection System (Invitrogen, Waltham, MA). Briefly, 1 x 10^5^ iPSCs in 10 µL Resuspension Buffer R were transfected with 10 pmol of SpyFi Cas9 Nuclease (Aldevron, Cat # 9214) and 50 pmol of the sgRNA. For siRNA treatment, 200 pmol siRNAs (Integrated DNA Technologies) were added to 1 x 10^5^ iPSCs (Table S6). The electroporation protocol for iPSCs was pulse voltage=1,100 V, pulse width=20 ms, pulse number=2. After electroporation, iPSCs were seeded into a well of Matrigel-coated 12-well plates in StemFlex medium (Gibco Cat # A3349401) supplemented with 10 µM Y-27632. After 2 days of electroporation, cells were detached with Accutase™ (StemCell Technologies, Cat # 07920) and followed by genomic DNA purification. For generating single-iPSC clones, *Sp*Cas9-sgRNA RNP complexes were delivered into iPSCs using the 4D-Nucleofector^TM^ electroporation system (Lonza, Basel, Switzerland) using Program CA-137. Briefly, 3 x 10^5^ iPSCs (C7) in 20 µL P3 primary cell solution (Lonza, Cat # V4XP-3032) were nucleofected with 40 pmol of SpyFi Cas9 Nuclease (Aldevron, Cat # 9214) and 200 pmol of the sgRNA. Immediately following nucleofection, the cells were seeded at low density onto Matrigel-coated 10-cm plates in StemFlex medium (Gibco Cat # A3349401) supplemented with 10 µM Y-27632 (StemCell Technologies Cat # 72302). Clonal colonies were manually picked 10 days after nucleofection, and re-adapted to mTeSR1 medium for expansion and routine maintenance.

### Genomic PCR for Sanger Sequencing and Amplicon NGS

Genomic DNA was purified by using the PureLink™ Genomic DNA kit (Invitrogen, Cat # K182002) according to the manufacturer’s protocol. For siRNA treated iPSCs, AllPrep DNA/RNA Micro Kit (Qiagen, Cat #80284) was used to purify both genomic DNA and RNA. For PCR amplification of the genomic region flanking *ATAD3A* exon 15, and *ATAD3B* exon 15, we designed primers via UCSC in silico PCR (https://genome.ucsc.edu/cgi-bin/hgPcr, Table S5). For Next Generation Sequencing, partial Illumina® adaptor sequences were added to the primers (Table S5). PCR reactions were performed using Q5® High-Fidelity DNA Polymerase (NEB Cat # M0491L). PCR products were purified with QIAquick PCR Purification kit (QIAGEN Cat# 28106) or QIAquick Gel Extraction Kit (QIAGEN Cat # 28706). Sanger sequencing of the PCR products was performed using Azenta Life Science (Burlington Massachusetts, USA). ICE (Inference of CRISPR Edits) analyses were used to determine HDR and indels frequencies. (Synthego Performance Analysis, ICE Analysis. 2019. v2.0. Synthego; [6.11.2021]). To analyze the contribution of gene-corrected sequences from Cas9-RNP treated iPSCs, the Sanger chromatogram of the same locus from SC20 was used as a control chromatogram, and c.1582C>T mock sequences were provided as donor templates on ICE analysis. Amplicon NGS was performed using the Amplicon-EZ service from Azenta Life Science.

### Whole Genome Sequencing and Genomic Analysis

Genomic DNA was purified from iPSCs using PureLink Genomic DNA Mini Kit (ThermoFisher Cat# K182002) according to the manufacture’s protocol. WGS library preparation was performed using TruSeq DNA PCR-free (550bp). Sequencing was performed on NovaSeq6000 S4 150PE to target 30x mappable (100Gb/sample). We aligned FASTQ files to the human reference genome GRCh37d5 (ftp://ftp-trace.ncbi.nih.gov/1000genomes/ftp/technical/reference/phase2_reference_assembly_sequence/hs37d5.fa.gz) using BWA-MEM (version 0.7.12). Duplicate reads were removed using the mark duplicate command of PICARD (version 1.130). Indel realignment and base quality recalibration were performed using GATK3 (version 2015.1-3.4.0-1-ga5ca3fc), resulting in analysis-ready BAM files. IGV screenshots were taken using Integrative Genomics Viewer (version 2.8.2). Read depth Manhattan plots for copy number profile were generated using CNVpytor (Suvakov et al. 2021).

Sequence similarities between the *ATAD3A* target region and its equivalent regions in two paralogs, *ATAD3B* and *ATAD3C*, were obtained by Smith-Waterman pairwise sequence alignment using the EBI EMBOSS Water web server (https://www.ebi.ac.uk/Tools/psa/emboss_water). For input sequences, we took the 320bp upstream and downstream sequence (641 bps total length) proximal to the target site of *ATAD3A* and the equivalent ones of *ATAD3B* and *ATAD3C*.

### Amplicon Sequencing Data Analysis

The amplicon sequencing paired reads were merged to single reads using PEAR (Zhang et al. 2014). Subsequently, the merged reads were aligned to GRCh37d5 using BWA-MEM and sorted by samtools (version 1.9). All the reads that are misaligned with the left and right positions of the genomic region targeted by the primer pair were filtered out. Allelic information for the target sites was calculated using samtools mpileup with the minimum base quality set to 20 and the minimum mapping quality set to 20 (Table S1 and S2).

### Interlocus Chimeric Read Discovery

The *ATAD3A* and *ATAD3B* amplicon sequences were queried against the human reference genome (GRCh37/hg19) using BLAT (UCSC Genome Browser) with default parameters. The alignments revealed an *ATAD3B*-specific sequence (TCatggtgtggggtccgcggccttgGCTGCCTCA) mapped to the *ATAD3A* amplicon with multiple gaps and an *ATAD3A*-specific sequence (TCgtgttgtgggagctGCTGCCTtggccggccCA) mapped to the *ATAD3B* amplicon, also with multiple gaps. These paralog-specific sequences were used as distinguishing markers for interlocus chimeric reads in the amplicon sequencing data. Identification of interlocus chimeric reads was performed using an in-house Bash script incorporating samtools view and grep, and boxplots were generated in Jupyter Notebook using the NumPy, Matplotlib, Pandas, and Seaborn packages.

### Long-read Amplicon Data Analysis

Long-range PCR was performed to amplify a 2,740 bp fragment (chr1:1,462,967-1,465,706). The resulting amplicon was sequenced using Oxford Nanopore Technology (ONT). Reads were aligned to the human reference genome (GRCh37d5) with Winnowmap (Jain et al. 2022). Four SNPs were located near the target mutation site: chr1:1,463,318C>A, chr1:1,463,337C>T, chr1:1,465,382A>G, and chr1:1,465,610G>A. These SNPs were used for haplotype genotyping except for chr1:1,463,337C>T, which has an adjacent 18-bp T homopolymer causing frequent indel errors in ONT sequencing. ONT reads spanning the target mutation and the selected SNPs were extracted, yielding 2,544 and 2,859 reads from iPSCs before and after Cas9-RNP treatment, respectively. Haplotypes were defined for each read based on the allelic combination of the target site and the three selected nearby SNPs (Figure 6A), resulting in 49 and 56 distinct haplotypes in the two iPSC samples.

Empirical sequencing error rates for each base (A, T, G, C) and deletion were estimated from mismatch frequencies across genomic positions of the amplicon, excluding the SNP positions and ± 100 bp around the Cas9-RNP target site. For all positions where reference base was 𝑏 ∈ {𝐴, 𝑇, 𝐺, 𝐶}, the empirical base error rate was defined as

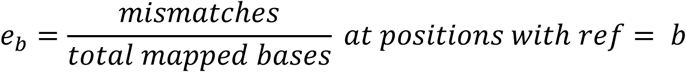

Similarly, the empirical deletion rate was defined as

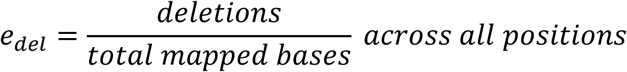

For a haplotype *h* spanning four informative positions (the target site and the selected three SNPs), the expected haplotype error probability was

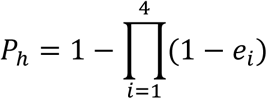

where *e_i_* is the error rate corresponding to the allele at position *i*.

The statistical significance of each haplotype was evaluated using a one-sided binomial test. Under the null hypothesis that haplotype-supporting reads arise solely from errors, the probability of observing at least *x* supporting reads was

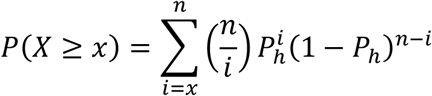

where *n* is the total number of reads. P-values were adjusted for multiple testing using the Bonferroni method, and haplotypes were considered significant if the adjusted p-value was < 0.01 (Table S3).

### Measurement of Mitochondrial Function

The oxygen consumption rate (OCR) was measured using Seahorse XFe24 analyzer by following the manufacturer’s protocol. Briefly, patient-derived and gene-corrected iPSCs were seeded at a density of 6 x 10^4^ cells per each well of Matrigel-coated XFe24 cell culture plates (Agilent 100777-004) a day before the measurement. The next day, cells were pre-incubated for an hour with complete XF DMEM medium (Agilent 103575-100) containing 10 mM glucose, 1 mM Sodium Pyruvate, and 2mM L-Glutamine. Electron Transport Chain (ETC) inhibitors were used by following working concentration: 1 μM Oligomycin, 0.5 μM Carbonyl cyanide 4-(trifluoromethoxy) phenylhydrazone (FCCP), 1 μM Antimycin A. After the measurement, each well of cells was lysed with 10 μL RIPA buffer. The protein concentration measured with the Bradford assay. Raw data were normalized with protein concentration in the Agilent program and analyzed in Microsoft Excel.

### Statistical Analysis

The GraphPad Prism and Excel software were used to process data, calculate statistics, and prepare graphs. The unpaired t-tests were used to determine statistical significance, with data presented as mean±SEM.

## Supporting information

Supplementary Information_ATAD3A_gene conversion

## LIST OF SUPPLEMENTARY MATERIALS

Supplementary Materials and Methods

Fig. S1. Karyotyping of patient-derived iPSCs carrying the heterozygous *ATAD3A* c.1582C>T variant

Fig. S2. Flow cytometry analysis of patient-derived iPSCs carrying the heterozygous *ATAD3A* c.1582C>T variant

Fig. S3. Alignment of the genomic sequences of the *ATAD3A* exon 15 and its homologous regions in *ATAD3B* and *ATAD3C*

Fig. S4. Sanger sequencing and ICE analysis of *ATAD3B* after delivering *ATAD3A* c.1582C>T variant-specific Cas9-RNP

Fig. S5. Sanger chromatograms of twenty subclones derived from C7 iPSCs electroporated with variant-specific Cas9-RNP

Fig. S6. High-resolution CNV plot for SC14 iPSC line.

Table S1. Allele frequencies of *ATAD3A* c.1582

Table S2. Allele frequencies of *ATAD3B* c.1582

Table S3. Haplotypes identified by read-backed phasing of ONT reads using SNP sites near the ATAD3A c.1582C>T variant site

Table S4. List of single-guided RNAs and corresponding sequences

Table S5. Primers utilized for the amplification of *ATAD3A* and *ATAD3B*

Table S6. List of siRNAs and their corresponding sequences

Table S7. List of primers used for qRT-PCR

## ACKNOWLEDGEMENTS

We thank the families for their participation in this study. We thank for the OMRF Center for Biomedical Data Sciences, specifically to Drs. Christopher Bottoms and David Stanford.

## AUTHOR CONTRIBUTIONS

Conceptualization: WHY

Methodology: TB, WHY, YP, JJK, MS, AA

Investigation: YP, TB, WHY, JJK, MS, AL, PZ, HP

Visualization: YP, TJ, WHY

Funding acquisition: WHY, AA, HVR, JRL, TH, JJK

Project administration: WHY

Supervision: WHY, AA

Writing – original draft: WHY

Writing – review & editing: WHY, TB, YP, AA, JJK, JRL

## DATA AND MATERIALS AVAILABILITY

All data associate with this study are present in the manuscript or the supplementary Materials.

## REFERENCES

Bai Y, Symington LS. 1996. A Rad52 homolog is required for RAD51-independent mitotic recombination in Saccharomyces cerevisiae. Genes Dev 10: 2025–2037.

Burnight ER, Gupta M, Wiley LA, Anfinson KR, Tran A, Triboulet R, Hoffmann JM, Klaahsen DL, Andorf JL, Jiao C et al. 2017. Using CRISPR-Cas9 to Generate Gene-Corrected Autologous iPSCs for the Treatment of Inherited Retinal Degeneration. Mol Ther 25: 1999–2013.

Carvalho CM, Lupski JR. 2016. Mechanisms underlying structural variant formation in genomic disorders. Nat Rev Genet 17: 224–238.

Chen JM, Cooper DN, Chuzhanova N, Ferec C, Patrinos GP. 2007. Gene conversion: mechanisms, evolution and human disease. Nat Rev Genet 8: 762–775.

Conant D, Hsiau T, Rossi N, Oki J, Maures T, Waite K, Yang J, Joshi S, Kelso R, Holden K et al. 2022. Inference of CRISPR Edits from Sanger Trace Data. CRISPR J 5: 123–130.

Cooper HM, Yang Y, Ylikallio E, Khairullin R, Woldegebriel R, Lin KL, Euro L, Palin E, Wolf A, Trokovic R et al. 2017. ATPase-deficient mitochondrial inner membrane protein ATAD3A disturbs mitochondrial dynamics in dominant hereditary spastic paraplegia. Hum Mol Genet 26: 1432–1443.

Desai R, Frazier AE, Durigon R, Patel H, Jones AW, Dalla Rosa I, Lake NJ, Compton AG, Mountford HS, Tucker EJ et al. 2017. *ATAD3* gene cluster deletions cause cerebellar dysfunction associated with altered mitochondrial DNA and cholesterol metabolism. Brain 140: 1595–1610.

Dilley RL, Verma P, Cho NW, Winters HD, Wondisford AR, Greenberg RA. 2016. Break-induced telomere synthesis underlies alternative telomere maintenance. Nature 539: 54–58.

Dorison N, Gaignard P, Bayot A, Gelot A, Becker PH, Fourati S, Lebigot E, Charles P, Wai T, Therond P et al. 2020. Mitochondrial dysfunction caused by novel *ATAD3A* mutations. Mol Genet Metab 131: 107–113.

Doudna JA, Charpentier E. 2014. Genome editing. The new frontier of genome engineering with CRISPR-Cas9. Science 346: 1258096.

Duret L, Galtier N. 2009. Biased gene conversion and the evolution of mammalian genomic landscapes. Annu Rev Genomics Hum Genet 10: 285–311.

Frazier AE, Compton AG, Kishita Y, Hock DH, Welch AE, Amarasekera SSC, Rius R, Formosa LE, Imai-Okazaki A, Francis D et al. 2021. Fatal perinatal mitochondrial cardiac failure caused by recurrent de novo duplications in the *ATAD3* locus. Med (N Y) 2: 49–73.

Gallagher DN, Pham N, Tsai AM, Janto NV, Choi J, Ira G, Haber JE. 2020. A Rad51-independent pathway promotes single-strand template repair in gene editing. PLoS Genet 16: e1008689.

Giannelli SG, Luoni M, Castoldi V, Massimino L, Cabassi T, Angeloni D, Demontis GC, Leocani L, Andreazzoli M, Broccoli V. 2018. Cas9/sgRNA selective targeting of the *P23H Rhodopsin* mutant allele for treating retinitis pigmentosa by intravitreal AAV9.PHP.B-based delivery. Hum Mol Genet 27: 761–779.

Gilquin B, Taillebourg E, Cherradi N, Hubstenberger A, Gay O, Merle N, Assard N, Fauvarque MO, Tomohiro S, Kuge O et al. 2010. The AAA+ ATPase ATAD3A controls mitochondrial dynamics at the interface of the inner and outer membranes. Mol Cell Biol 30: 1984–1996.

Gunning AC, Strucinska K, Munoz Oreja M, Parrish A, Caswell R, Stals KL, Durigon R, Durlacher-Betzer K, Cunningham MH, Grochowski CM et al. 2020. Recurrent De Novo NAHR Reciprocal Duplications in the *ATAD3* Gene Cluster Cause a Neurogenetic Trait with Perturbed Cholesterol and Mitochondrial Metabolism. Am J Hum Genet 106: 272–279.

Gyorgy B, Nist-Lund C, Pan B, Asai Y, Karavitaki KD, Kleinstiver BP, Garcia SP, Zaborowski MP, Solanes P, Spataro S et al. 2019. Allele-specific gene editing prevents deafness in a model of dominant progressive hearing loss. Nat Med 25: 1123–1130.

Hanes I, McMillan HJ, Ito Y, Kernohan KD, Lazier J, Lines MA, Dyment DA. 2020. A splice variant in *ATAD3A* expands the clinical and genetic spectrum of Harel-Yoon syndrome. Neurol Genet 6: e452.

Harel T, Yoon WH, Garone C, Gu S, Coban-Akdemir Z, Eldomery MK, Posey JE, Jhangiani SN, Rosenfeld JA, Cho MT et al. 2016. Recurrent De Novo and Biallelic Variation of *ATAD3A*, Encoding a Mitochondrial Membrane Protein, Results in Distinct Neurological Syndromes. Am J Hum Genet 99: 831–845.

He J, Mao CC, Reyes A, Sembongi H, Di Re M, Granycome C, Clippingdale AB, Fearnley IM, Harbour M, Robinson AJ et al. 2007. The AAA+ protein ATAD3 has displacement loop binding properties and is involved in mitochondrial nucleoid organization. J Cell Biol 176: 141–146.

Hsu PD, Scott DA, Weinstein JA, Ran FA, Konermann S, Agarwala V, Li Y, Fine EJ, Wu X, Shalem O et al. 2013. DNA targeting specificity of RNA-guided Cas9 nucleases. Nat Biotechnol 31: 827–832.

Inoue K, Lupski JR. 2002. Molecular mechanisms for genomic disorders. Annu Rev Genomics Hum Genet 3: 199–242.

Ira G, Haber JE. 2002. Characterization of *RAD51*-independent break-induced replication that acts preferentially with short homologous sequences. Mol Cell Biol 22: 6384–6392.

Jackson SP, Bartek J. 2009. The DNA-damage response in human biology and disease. Nature 461: 1071–1078.

Jain C, Rhie A, Hansen NF, Koren S, Phillippy AM. 2022. Long-read mapping to repetitive reference sequences using Winnowmap2. Nat Methods 19: 705–710.

Javidi-Parsijani P, Lyu P, Makani V, Sarhan WM, Yoo KW, El-Korashi L, Atala A, Lu B. 2020. CRISPR/Cas9 increases mitotic gene conversion in human cells. Gene Ther 27: 281–296.

Jin G, Xu C, Zhang X, Long J, Rezaeian AH, Liu C, Furth ME, Kridel S, Pasche B, Bian XW et al. 2018. Atad3a suppresses Pink1-dependent mitophagy to maintain homeostasis of hematopoietic progenitor cells. Nat Immunol 19: 29–40.

Jinek M, Chylinski K, Fonfara I, Hauer M, Doudna JA, Charpentier E. 2012. A programmable dual-RNA-guided DNA endonuclease in adaptive bacterial immunity. Science 337: 816–821.

Jonkman MF, Pasmooij AM. 2009. Revertant mosaicism--patchwork in the skin. N Engl J Med 360: 1680–1682.

Jonkman MF, Scheffer H, Stulp R, Pas HH, Nijenhuis M, Heeres K, Owaribe K, Pulkkinen L, Uitto J. 1997. Revertant mosaicism in epidermolysis bullosa caused by mitotic gene conversion. Cell 88: 543–551.

Kim S, Kim D, Cho SW, Kim J, Kim JS. 2014. Highly efficient RNA-guided genome editing in human cells via delivery of purified Cas9 ribonucleoproteins. Genome Res 24: 1012–1019.

Koike-Yusa H, Li Y, Tan EP, Velasco-Herrera Mdel C, Yusa K. 2014. Genome-wide recessive genetic screening in mammalian cells with a lentiviral CRISPR-guide RNA library. Nat Biotechnol 32: 267–273.

Kosicki M, Tomberg K, Bradley A. 2018. Repair of double-strand breaks induced by CRISPR-Cas9 leads to large deletions and complex rearrangements. Nat Biotechnol 36: 765–771.

LaRocque JR, Stark JM, Oh J, Bojilova E, Yusa K, Horie K, Takeda J, Jasin M. 2011. Interhomolog recombination and loss of heterozygosity in wild-type and *Bloom syndrome helicase* (*BLM*)-deficient mammalian cells. Proc Natl Acad Sci U S A 108: 11971–11976.

Li S, Lamarche F, Charton R, Delphin C, Gires O, Hubstenberger A, Schlattner U, Rousseau D. 2014. Expression analysis of *ATAD3* isoforms in rodent and human cell lines and tissues. Gene 535: 60–69.

Liang X, Potter J, Kumar S, Zou Y, Quintanilla R, Sridharan M, Carte J, Chen W, Roark N, Ranganathan S et al. 2015. Rapid and highly efficient mammalian cell engineering via Cas9 protein transfection. J Biotechnol 208: 44–53.

Lieber MR. 2010. The mechanism of double-strand DNA break repair by the nonhomologous DNA end-joining pathway. Annu Rev Biochem 79: 181–211.

Ma H, Marti-Gutierrez N, Park SW, Wu J, Hayama T, Darby H, Van Dyken C, Li Y, Koski A, Liang D et al. 2018. Ma et al. reply. Nature 560: E10–E23.

Ma H, Marti-Gutierrez N, Park SW, Wu J, Lee Y, Suzuki K, Koski A, Ji D, Hayama T, Ahmed R et al. 2017. Correction of a pathogenic gene mutation in human embryos. Nature 548: 413–419.

Mayle R, Campbell IM, Beck CR, Yu Y, Wilson M, Shaw CA, Bjergbaek L, Lupski JR, Ira G. 2015. DNA REPAIR. Mus81 and converging forks limit the mutagenicity of replication fork breakage. Science 349: 742–747.

Peralta S, Gonzalez-Quintana A, Ybarra M, Delmiro A, Perez-Perez R, Docampo J, Arenas J, Blazquez A, Ugalde C, Martin MA. 2019. Novel *ATAD3A* recessive mutation associated to fatal cerebellar hypoplasia with multiorgan involvement and mitochondrial structural abnormalities. Mol Genet Metab 128: 452–462.

Pharaoh G, Sataranatarajan K, Street K, Hill S, Gregston J, Ahn B, Kinter C, Kinter M, Van Remmen H. 2019. Metabolic and Stress Response Changes Precede Disease Onset in the Spinal Cord of Mutant *SOD1* ALS Mice. Front Neurosci 13: 487.

Prakash R, Zhang Y, Feng W, Jasin M. 2015. Homologous recombination and human health: the roles of BRCA1, BRCA2, and associated proteins. Cold Spring Harb Perspect Biol 7: a016600.

Rabai A, Reisser L, Reina-San-Martin B, Mamchaoui K, Cowling BS, Nicot AS, Laporte J. 2019. Allele-Specific CRISPR/Cas9 Correction of a Heterozygous *DNM2* Mutation Rescues Centronuclear Myopathy Cell Phenotypes. Mol Ther Nucleic Acids 16: 246–256.

Richardson CD, Kazane KR, Feng SJ, Zelin E, Bray NL, Schafer AJ, Floor SN, Corn JE. 2018. CRISPR-Cas9 genome editing in human cells occurs via the Fanconi anemia pathway. Nat Genet 50: 1132–1139.

Saini N, Ramakrishnan S, Elango R, Ayyar S, Zhang Y, Deem A, Ira G, Haber JE, Lobachev KS, Malkova A. 2013. Migrating bubble during break-induced replication drives conservative DNA synthesis. Nature 502: 389–392.

Smith C, Abalde-Atristain L, He C, Brodsky BR, Braunstein EM, Chaudhari P, Jang YY, Cheng L, Ye Z. 2015. Efficient and allele-specific genome editing of disease loci in human iPSCs. Mol Ther 23: 570–577.

Suvakov M, Panda A, Diesh C, Holmes I, Abyzov A. 2021. CNVpytor: a tool for copy number variation detection and analysis from read depth and allele imbalance in whole-genome sequencing. Gigascience 10.

Sy SM, Huen MS, Chen J. 2009. PALB2 is an integral component of the BRCA complex required for homologous recombination repair. Proc Natl Acad Sci U S A 106: 7155–7160.

Tan EP, Li Y, Velasco-Herrera Mdel C, Yusa K, Bradley A. 2015. Off-target assessment of CRISPR-Cas9 guiding RNAs in human iPS and mouse ES cells. Genesis 53: 225–236.

van Overbeek M, Capurso D, Carter MM, Thompson MS, Frias E, Russ C, Reece-Hoyes JS, Nye C, Gradia S, Vidal B et al. 2016. DNA Repair Profiling Reveals Nonrandom Outcomes at Cas9-Mediated Breaks. Mol Cell 63: 633–646.

Wong AK, Ormonde PA, Pero R, Chen Y, Lian L, Salada G, Berry S, Lawrence Q, Dayananth P, Ha P et al. 1998. Characterization of a carboxy-terminal BRCA1 interacting protein. Oncogene 17: 2279–2285.

Xia B, Dorsman JC, Ameziane N, de Vries Y, Rooimans MA, Sheng Q, Pals G, Errami A, Gluckman E, Llera J et al. 2007. Fanconi anemia is associated with a defect in the BRCA2 partner PALB2. Nat Genet 39: 159–161.

Yamamoto Y, Makiyama T, Harita T, Sasaki K, Wuriyanghai Y, Hayano M, Nishiuchi S, Kohjitani H, Hirose S, Chen J et al. 2017. Allele-specific ablation rescues electrophysiological abnormalities in a human iPS cell model of long-QT syndrome with a CALM2 mutation. Hum Mol Genet 26: 1670–1677.

Yanovsky-Dagan S, Frumkin A, Lupski JR, Harel T. 2022. CRISPR/Cas9-induced gene conversion between *ATAD3* paralogs. HGG Adv 3: 100092.

Yap ZY, Park YH, Wortmann SB, Gunning AC, Ezer S, Lee S, Duraine L, Wilichowski E, Wilson K, Mayr JA et al. 2021. Functional interpretation of ATAD3A variants in neuro-mitochondrial phenotypes. Genome Med 13: 55.

Yu X, Wu LC, Bowcock AM, Aronheim A, Baer R. 1998. The C-terminal (BRCT) domains of BRCA1 interact in vivo with CtIP, a protein implicated in the CtBP pathway of transcriptional repression. J Biol Chem 273: 25388–25392.

Zhang J, Kobert K, Flouri T, Stamatakis A. 2014. PEAR: a fast and accurate Illumina Paired-End reAd mergeR. Bioinformatics 30: 614–620.

