## Supplementary Information_ATAD3A_gene conversion for "Allele-specific correction of *ATAD3A* pathogenic variants via template-free CRISPR-Cas9 editing and gene conversion"

### SUPPLEMENTARY MATERIALS AND METHODS

#### Flow cytometry

iPS cells grown in 6-well plates were harvested using 0.05% Trypsin-EDTA and stained for cells surface antigens using combinations of the following antibodies: IgG1 Alexa 488 negative control (AbD Serotec MCA2356A488), 1:10; anti-CD29 Alexa 488 (AbD Serotec MCA2298A488), 1:10; or IgG3 anti-SSEA4-APC (R&D FAB1435A), 1:10. Cells then fixed with 2% paraformaldehyde, and permeabilized with 0.1% Saponin and 0.1% BSA in DPBS. Nuclear antigens were stained with mouse IgG1-PE negative control (BD 559320), 1:10; or mouse IgG1 anti-OCT4-PE (BD 560186), 1:10. Stained cells were detected by cell cytometry (BD LSRII Analyzer) and the data was analyzed with BD FACS Diva software.

#### iPSCs Karyotyping

For C7, C10, C27, and C30 iPSC lines, karyotyping by G-banding was performed by Baylor Genetics (Houston, Texas, USA). For TM-C9 and TM-C21 iPSC lines, KaryoStat™ assay was performed by Thermo Fisher Scientific (Carlsbad, California, USA).

#### Quantitative real-time RT-PCR

Total RNA from iPSCs was extracted using the AllPrep® DNA/RNA Micro Kit (Qiagen, Cat #80284), followed by cDNA synthesis using iScript cDNA synthesis kit (Bio-Rad #1708891). Quantitative RT-PCR was performed using the FastStart Essential DNA Green Master (Roche #6402712001) and LightCycler® 96 Instrument (Roche). Amplification signals were normalized to *GAPDH*, and fold-changes were calculated using ΔΔCt method. Data analysis and calculations were performed using Excel (Microsoft). Primers used for qRT-PCR are in Table S7.

SUPPLEMENTARY FIGURES

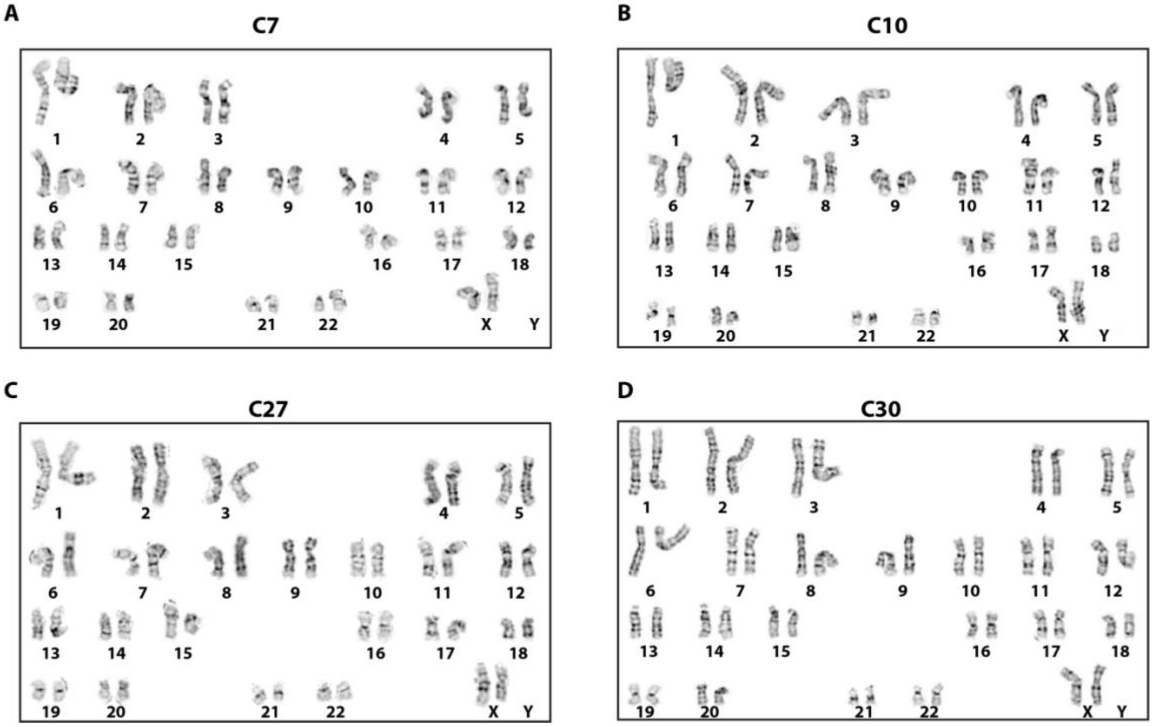


**Fig. S1. Karyotyping of patient-derived iPSCs carrying the heterozygous *ATAD3A* c.1582C>T variant**

**(A-B)** Fibroblasts obtained from the first individual carrying the *de novo ATAD3A* c.1582C>T variant were reprogrammed into two iPSC clones: C7 and C10. Both C7 and C10 iPSC lines exhibit normal 46, XX karyotypes. **(C-D)** iPSC clones C27 and C30 were generated from PBMCs of a second individual carrying the identical heterozygous *ATAD3A* variant (c.1582C>T), and display normal 46, XX karyotypes.

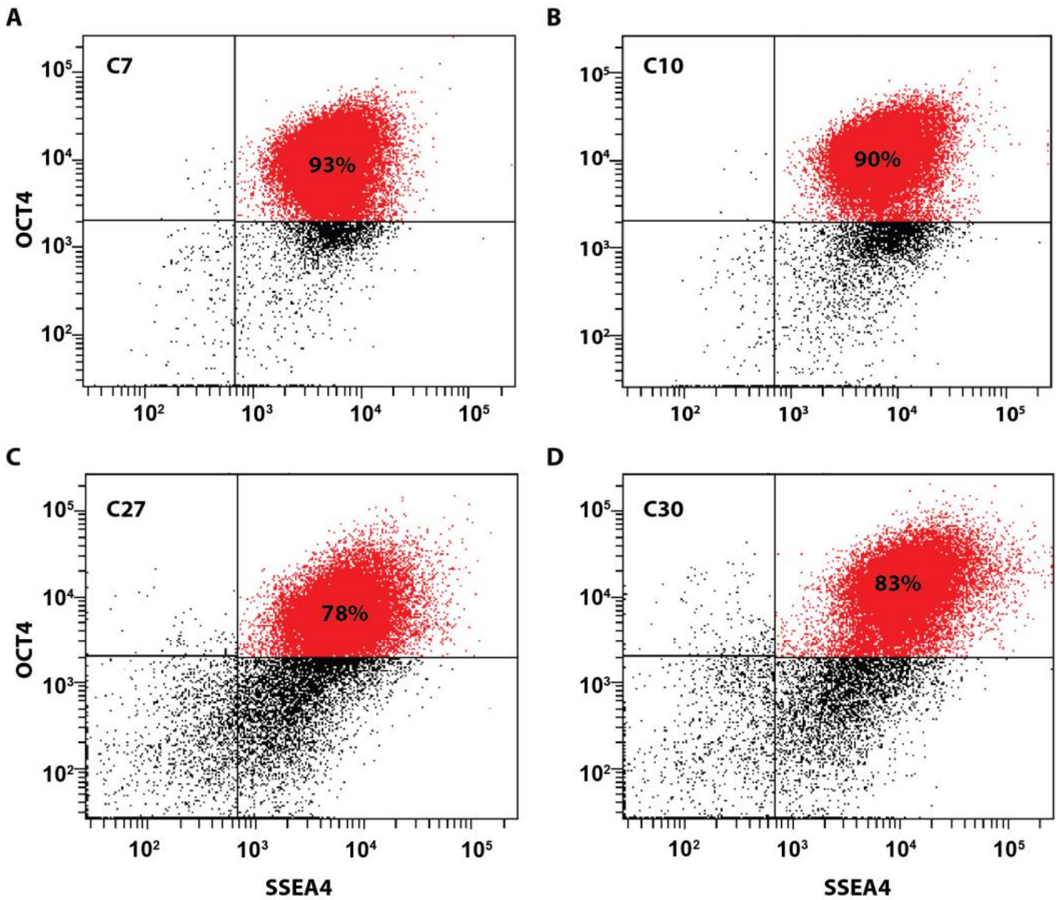


**Fig. S2. Flow cytometry analysis of patient-derived iPSCs carrying the heterozygous *ATAD3A* c.1582C>T variant**

Flow cytometry analysis of **(A)** C7, **(B)** C10, **(C)** C27, and **(D)** C30 iPSC lines using antibodies for OCT4, and SSEA4, hallmarks of pluripotent stem cells.

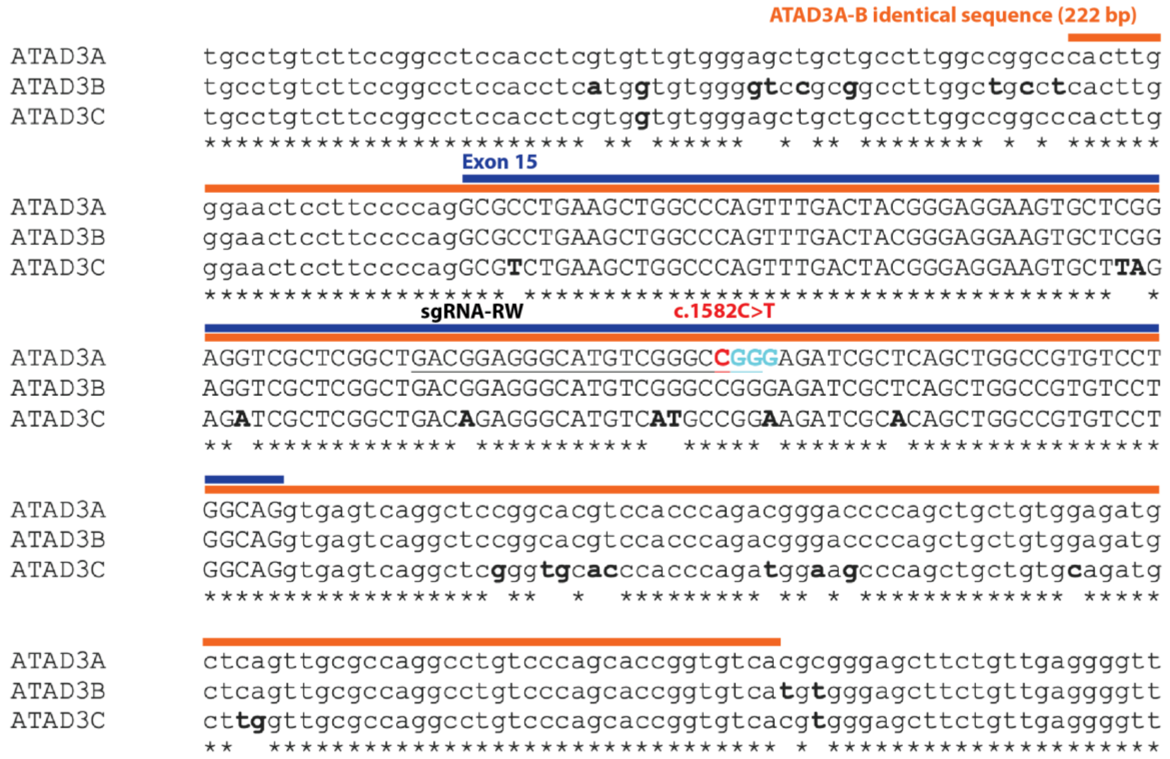


**Fig. S3. Alignment of the genomic sequences of the *ATAD3A* exon 15 and its homologous regions in *ATAD3B* and *ATAD3C***

Alignments of genomic sequences from *ATAD3A* exon 15, *ATAD3B* exon 15, and *ATAD3C* Exon 11 using the EMBL-EBI search and sequence analysis tool. The orange bar highlights a 222 bp sequence identical between *ATAD3A* and *ATAD3B*. The blue bar with uppercase sequences indicates exon 15 of *ATAD3A*. The c.1582 C allele in *ATAD3A* is marked in red. The underlined sequence shows the sgRNA-RW targeting site. Bold sequences in *ATAD3B* and *ATAD3C* display unique nucleotides in *ATAD3B* and *ATAD3C* that differ from *ATAD3A*.

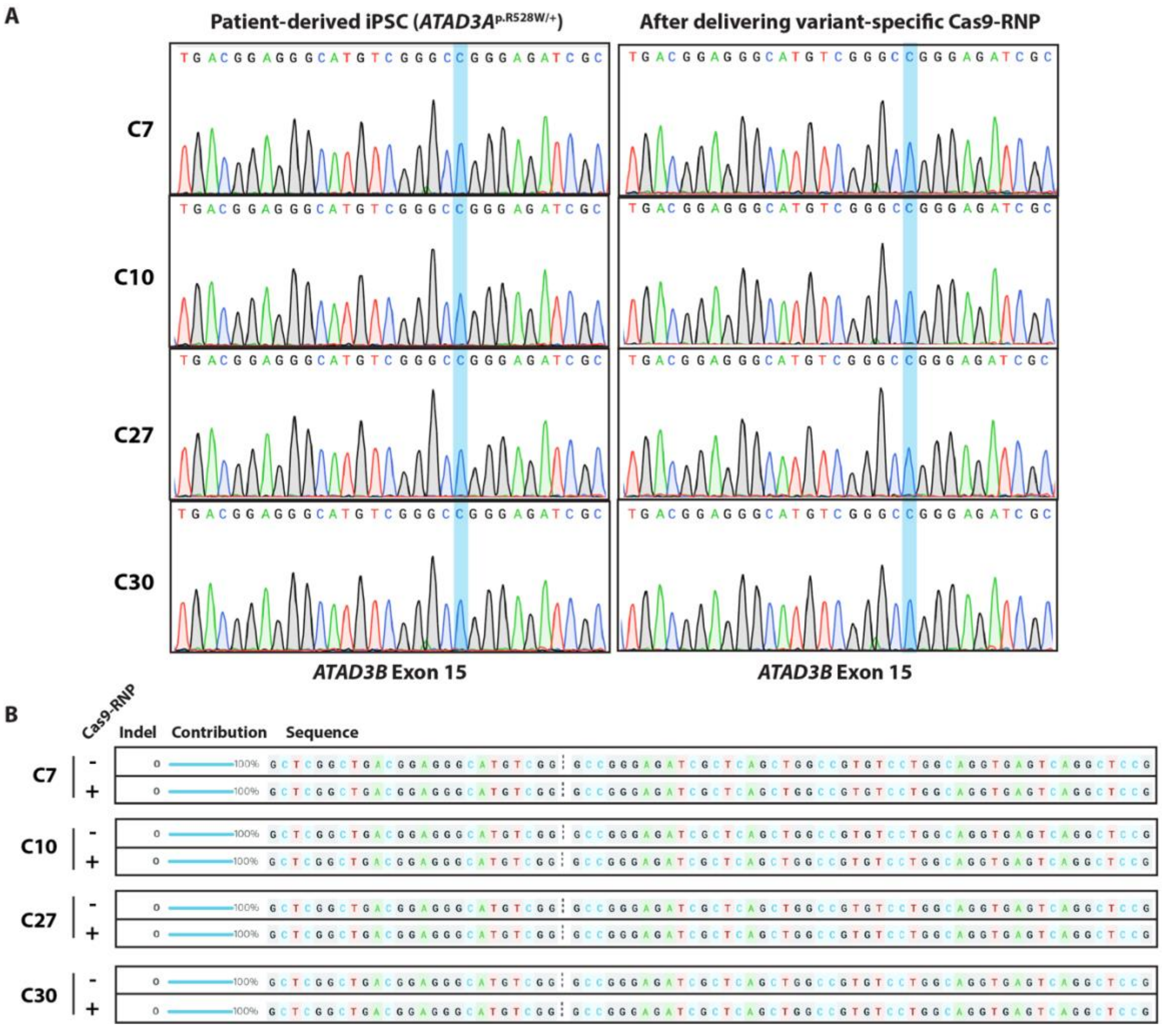


**Fig. S4. Sanger sequencing and ICE analysis of *ATAD3B* after delivering *ATAD3A* c.1582C>T variant-specific Cas9-RNP**

**(A)** Sanger chromatograms for *ATAD3B* exon 15 from C7, C10, C27, and C30 iPSC lines before (left) and after (right) delivery of the *ATAD3A* c.1582C>T variant-specific Cas9-RNP. Blue boxes highlight the *ATAD3B* c.1582C allele in exon 15, which is homologous to the *ATAD3A* c.1582C allele. **(B)** ICE analyses for the *ATAD3B* Sanger chromatograms reveal no indels in *ATAD3B* exon 15 both before and after the delivery of variant-specific Cas9-RNP.

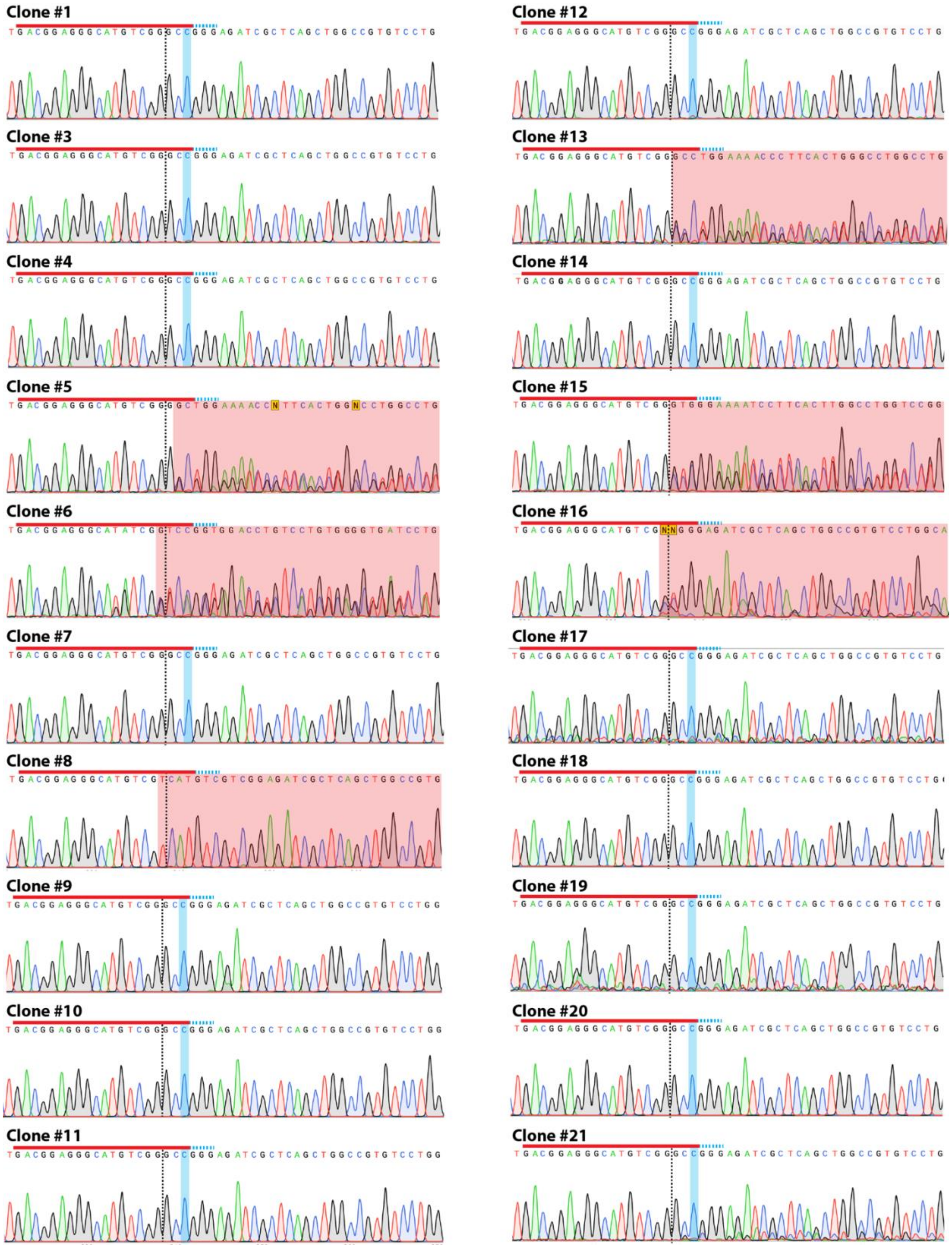
Clone #1
Clone #3
Clone #4
Clone #5
Clone #6
Clone #7
Clone #8
Clone #9
Clone #10
Clone #11
Clone #12
Clone #13
Clone #14
Clone #15
Clone #16
Clone #17
Clone #18
Clone #19
Clone #20
Clone #21

**Fig. S5. Sanger chromatograms of twenty subclones derived from C7 iPSCs electroporated with variant-specific Cas9-RNP**

Sanger chromatograms for the *ATAD3A* exon 15 regions of twenty subclones derived from C7 iPSCs following c.1582C>T variant -specific Cas9 RNP delivery. Red bars highlight the target sequences of the variant-specific sgRNA. Cyan dots indicate the PAM sequences. Black dotted lines indicate the Cas9 cutting site. Blue boxes indicate the correction (c.1582 C) of the *ATAD3A* c.1582C>T pathogenic variant in 14 subclones, whereas the other 6 subclones exhibit indels (indicated by red boxes).

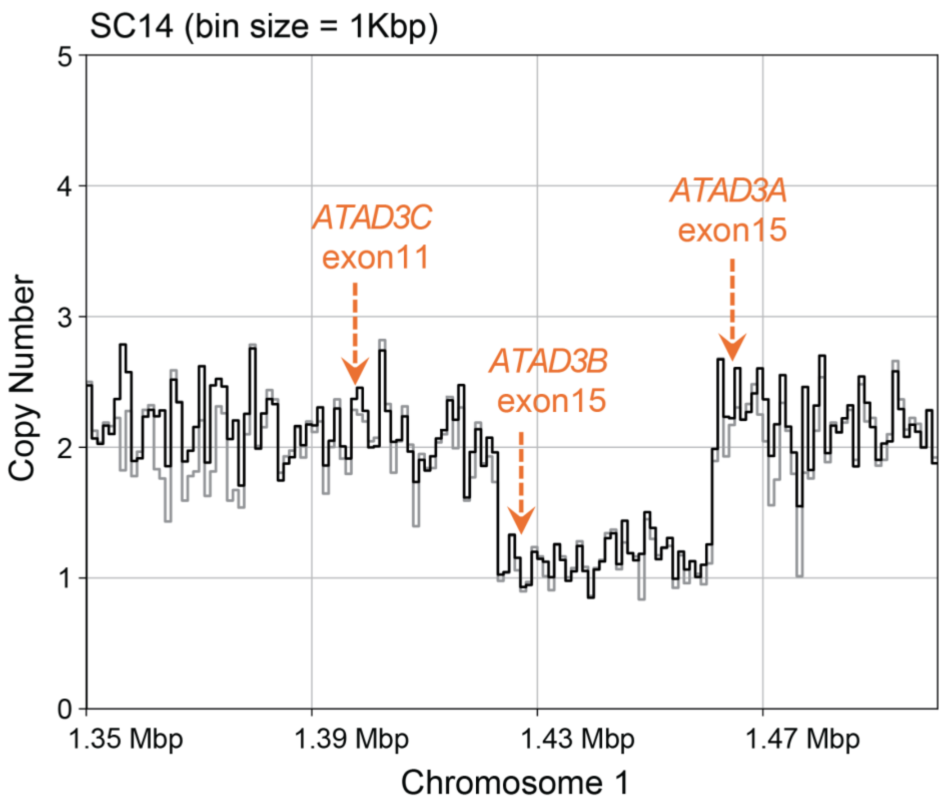


**Figure S6**

**Figure S6. High-resolution copy number profile confirming the deletion spanning ATAD3A and ATAD3B**

Copy number profile across chromosome 1 (bin size = 1 kb) showing read depth in the SC20 iPSC clone. A deletion is evident between the immediate upstream regions of *ATAD3B* exon 15 and *ATAD3A* exon 15. The breakpoints are located very close to these exons, which share highly similar sequences, suggesting that sequence homology may have mediated the rearrangement. The positions of *ATAD3C* exon 11, *ATAD3B* exon 15, and *ATAD3A* exon 15 are indicated.

### SUPPLEMENTARY TABLES

#### Table S1. Allele frequencies of *ATAD3A* c.1582

| iPSC lines | Cas9-RNP | C (intact) | T (intact) | C (indel) | T (indel) | Deletion | Other Bases | Corrected cell % |
|---|---|---|---|---|---|---|---|---|
| C7 | - | 50.1% | 49.0% | 0.4% | 0.5% | 0.1% | 0.1% | |
| | sgRNA-RW | 71.8% | 6.7% | 4.5% | 9.2% | 7.5% | 0.2% | 53.1% |
| C10 | - | 50.6% | 47.6% | 0.5% | 0.8% | 0.4% | 0.0% | |
| | sgRNA-RW | 66.4% | 13.3% | 2.6% | 9.7% | 7.8% | 0.3% | 38.6% |
| C27 | - | 50.1% | 49.0% | 0.4% | 0.4% | 0.1% | 0.1% | |
| | sgRNA-RW | 69.7% | 7.9% | 3.2% | 11.8% | 7.0% | 0.3% | 46.4% |
| C30 | - | 50.8% | 48.2% | 0.4% | 0.5% | 0.2% | 0.0% | |
| | sgRNA-RW | 65.2% | 12.2% | 3.3% | 9.7% | 9.2% | 0.3% | 37.7% |

#### Table S2. Allele frequencies of *ATAD3B* c.1582

| iPSC lines | Cas9-RNP | C (intact) | T (intact) | C (indel) | T (indel) | Deletion | Other Bases |
|---|---|---|---|---|---|---|---|
| C7 | - | 96.6% | 0.0% | 3.4% | 0.0% | 0.0% | 0.0% |
| | sgRNA-RW | 91.8% | 0.0% | 7.0% | 0.1% | 1.0% | 0.1% |
| C10 | - | 95.9% | 0.0% | 4.1% | 0.0% | 0.0% | 0.0% |
| | sgRNA-RW | 94.4% | 0.0% | 5.0% | 0.0% | 0.5% | 0.0% |
| C27 | - | 96.5% | 0.1% | 3.4% | 0.0% | 0.0% | 0.0% |
| | sgRNA-RW | 94.1% | 0.0% | 5.4% | 0.0% | 0.4% | 0.0% |
| C30 | - | 95.7% | 0.0% | 4.3% | 0.0% | 0.0% | 0.0% |
| | sgRNA-RW | 92.1% | 0.0% | 6.6% | 0.1% | 1.1% | 0.0% |

Table S3. Haplotypes identified by read-backed phasing of ONT reads using SNP sites near the *ADAD3A* c.1582C>T variant

| Haplotype | Control (S77p20inv) | | | | | | Cas9-RNP (S77p28inv-gR3) | | | | | |
|---|---|---|---|---|---|---|---|---|---|---|---|---|
| (-SNP1-Target-SNP2-SNP3-) | Read count | Read fraction | Haplotype error probability | *P*-value | Adjusted *P*-value (Bonferroni) | Significance | Read count | Read fraction | Haplotype error probability | *P*-value | Adjusted *P*-value (Bonferroni) | Significance |
| -C-C-A-G- | 1036 | 42.70% | 0.002153 | 0 | 0 | TRUE | 1263 | 46.64% | 0.002023 | 0 | 0 | TRUE |
| -A-T-G-A- | 1005 | 41.43% | 0.002120 | 0 | 0 | TRUE | 408 | 15.07% | 0.002247 | 0 | 0 | TRUE |
| -A-C-G-A- | 106 | 4.37% | 0.002003 | 6.78E-99 | 3.32E-97 | TRUE | 595 | 21.97% | 0.001886 | 0 | 0 | TRUE |
| -A-DEL-G-A- | 2 | 0.08% | 0.002796 | 0.9934 | 1 | FALSE | 229 | 8.46% | 0.002998 | 4.33E-237 | 2.43E-235 | TRUE |
| -A-C-A-G- | 104 | 4.29% | 0.002003 | 3.15E-96 | 1.54E-94 | TRUE | 93 | 3.43% | 0.001886 | 1.24E-79 | 6.95E-78 | TRUE |
| -C-T-G-A- | 72 | 2.97% | 0.002270 | 1.60E-52 | 7.82E-51 | TRUE | 30 | 1.11% | 0.002384 | 4.89E-11 | 2.74E-09 | TRUE |
| -C-C-G-A- | 43 | 1.77% | 0.002153 | 3.43E-24 | 1.68E-22 | TRUE | 73 | 2.70% | 0.002023 | 1.49E-53 | 8.35E-52 | TRUE |
| -A-T-A-G- | 33 | 1.36% | 0.002120 | 7.64E-16 | 3.75E-14 | TRUE | 5 | 0.18% | 0.002247 | 0.7682 | 1 | FALSE |
| -C-C-A-A- | 25 | 1.03% | 0.001249 | 9.90E-15 | 4.85E-13 | TRUE | 12 | 0.44% | 0.001307 | 0.0005 | 0.0283 | FALSE |

**Table S3. Haplotypes identified by read-backed phasing of ONT reads using SNP sites near the ADAD3A c.1582C>T variant site**

ONT reads spanning the target site and three nearby SNPs were tested for the presence of true haplotypes beyond the expected frequency from haplotype errors. Haplotype error probability was estimated from the empirical sequencing error rate in each dataset. For each haplotype, read count, fraction, estimated haplotype error probability, P-value, Bonferroni-adjusted P-value, and significance status are shown for control (untreated iPSCs) and Cas9-RNP–treated iPSCs. A binomial test using haplotype error probability and total read counts in each dataset was applied to evaluate significance (Methods).

**Table S4. List of single-guided RNAs and corresponding sequences**

| sgRNA | Sequences | PAM |
|---|---|---|
| sgRNA-RW | 5'-GACGGAGGGCAUGUCGGGC**U**-3' | GGG |

**Table S5. Primers utilized for the amplification of *ATAD3A* and *ATAD3B***

| Primer | Sequences | Target |
|---|---|---|
| ATAD3A-RW_F4 | 5'-CCTGCAGCCACTCCCTGCTC-3' | *ATAD3A* Exon15 |
| ATAD3A-RW_R4 | 5′-CCCTCAACAGAAGCTCCCGCG-3′ | |
| ATAD3A-RW-Illumina_F | 5'-**ACACTCTTTCCCTACACGACGCTCTTCCGATCT** CCTGCAGCCACTCCCTGCTC-3' | |
| ATAD3A-RW-Illumina_R | 5'-**GACTGGAGTTCAGACGTGTGCTCTTCCGATCT** CCCTCAACAGAAGCTCCCGCG-3' | |
| ATAD3A-long new_F4 | 5′-GTTTCCCGCCACTTTAGGT-3′ | |
| ATAD3A-long new_R1.5A | 5′-CCTGGCTTCAAATAATCTAACTTGG-3′ | |
| ATAD3B-RW_F1 | 5′-CCGGCCACAGAAGGAAAACGGTG-3′ | *ATAD3B* Exon15 |
| ATAD3B-RW_R3 | 5′-CCCTCAACAGAAGCTCCCACA-3′ | |
| ATAD3B-RW-Illumina_F | 5'-**ACACTCTTTCCCTACACGACGCTCTTCCGATCT** CCGGCCACAGAAGGAAAACGGTG-3' | |
| ATAD3B-RW-Illumina_R | 5'-**GACTGGAGTTCAGACGTGTGCTCTTCCGATCT** CCCTCAACAGAAGCTCCCACA-3' | |

**Table S6. List of siRNAs and their corresponding sequences**

| siRNA | Duplex Sequences |
|---|---|
| Scrambled | 5′- CGUUAAUCGCGUAUAAUACGCGUAT-3′<br>3′-CAGCAAUUAGCGCAUAUUAUGCGCAUA-5′ |
| siRAD51 | 5′- GUCACAAACUGAUCUAAAAUGUUTA-3′<br>3′-AUCAGUGUUUGACUAGAUUUUACAAAU-5′ |
| siBRCA1 | 5′- GUACGAGAUUUAGUCAACUUGUUGA-3′<br>3′-UUCAUGCUCUAAAUCAGUUGAACAACU-5′ |
| siBRCA2 | 5′- CAAGAAGCAUGUCAUGGUAAUACTT-3′<br>3′-GAGUUCUUCGUACAGUACCAUUAUGAA-5′ |
| siCtIP | 5′- GAGAAUGUUUUAGAUGACAUAAAGA-3′<br>3′-GACUCUUACAAAAUCUACUGUAUUUCU-5′ |

**Table S7. List of primers used for qRT-PCR**

| Primer for qRT-PCR | Sequences |
|---|---|
| RAD51-F | 5′-AGACCGAGCCCTAAGGAGAG-3′ |
| RAD51-R | 5′-CTTCTCTACTCGCTTGCCCC-3′ |
| CtIP-F | 5′-CAGAATAGGACTGAGTACGGTAAAG-3′ |
| CtIP-R | 5′-CTGACTGCCATCCTTTGTATCT-3′ |
| BRCA1-F | 5′-AAGCTGACAGATGGTTCATT-3′ |
| BRCA1-R | 5′-ACAGGTTCCTTGATCAACTC-3′ |
| BRCA2-F | 5′-TGTGGAAGTTGCGTATTGTA-3′ |
| BRCA2-R | 5′-TAAATCTGATGATGGACGCC-3′ |
